# Preorganized RdRp-Thumb Dynamics Drives SARS-CoV-2 Polymerase Function

**DOI:** 10.64898/2026.05.29.728794

**Authors:** Hannah Gibbs, Artiom Butuc, Preston B Moore, Eleonora Gianti

## Abstract

The SARS-CoV-2 RNA-dependent RNA polymerase drives viral genome replication and is a major antiviral target. How intrinsic conformational dynamics organize functional states of the polymerase, however, remains incompletely understood. Here, molecular dynamics (MD) simulations combined with free-energy landscape analysis reveal that the apo polymerase samples preexisting conformational states defined by coordinated thumb-subdomain motions. Projection of experimental structures, representative of the polymerase nucleic acid cycle, onto the conformational landscape identified discrete basins spanning apo-like and elongation-like states and revealed a coherent structural axis coupling global polymerase compaction (radius of gyration, Rg) with thumb–interface separation (center-of-mass distance, COM). These motions connect catalytic motifs, RNA-binding regions, and distal regulatory elements across the polymerase ensemble.

The observed conformational organization is not apparent from static structures alone and supports a model in which functional transitions arise from intrinsic collective dynamics of the apo enzyme. Intermediate conformational ensembles combine structural stability with retained inter-domain flexibility, identifying mechanically responsive states favorable for allosteric modulation. Together, these findings define structurally coupled regulatory regions within the coronavirus polymerase and support conformational trapping of thumb-subdomain dynamics as a potential strategy for antiviral design targeting RNA virus replication machinery.

## INTRODUCTION

The SARS-CoV-2 RNA-dependent RNA polymerase (RdRp) is the central engine of viral genome replication and transcription, making it a primary determinant of coronavirus (CoV) proliferation and a major antiviral target. High-resolution experimental structures have established the architecture of the replication and transcription complex (RTC) and clarified how RNA templates, nucleotides, and cofactors engage the enzyme during RNA synthesis.^1,2^ Like other viral polymerases, the enzyme adopts the canonical right-hand architecture (**Fig. 1**), composed of fingers, palm, and thumb subdomains that together form the catalytic cavity.^3^ However, static structures alone cannot resolve how conformational fluctuations organize functional transitions required for catalysis and RNA translocation.

**Figure 1.**
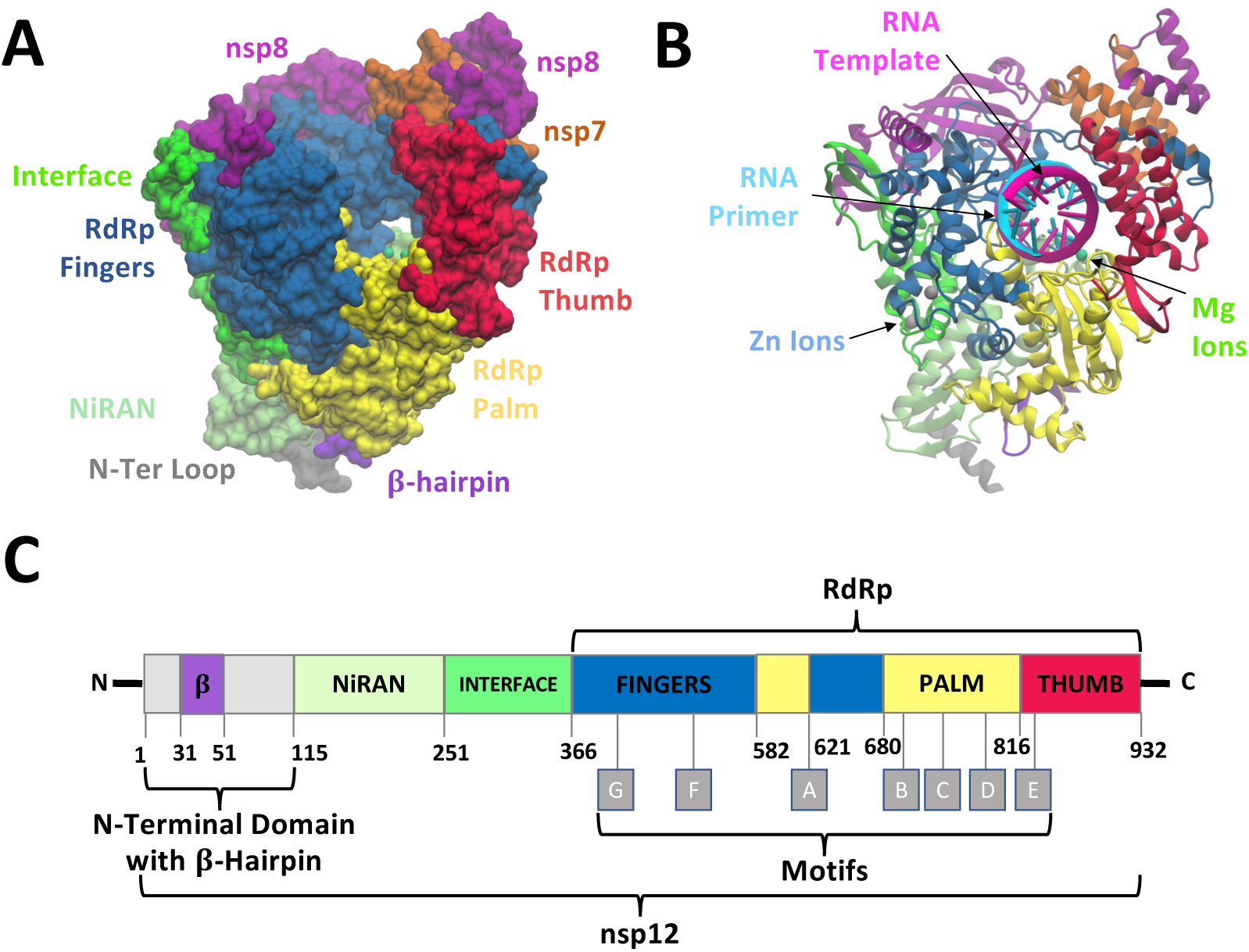
The SARS-CoV-2 core polymerase complex (nsp12–nsp8/nsp7). (A) Three-dimensional structure of nsp12 in complex with cofactors nsp8 (two molecules) and nsp7 (one molecule), forming the replication–transcription complex (RTC). The structure is shown in the apo state (PDB ID: 6M71). (B) Structure of RTC in complex with RNA (PDB ID: 7BZF). (C) Architecture of nsp12 domains, subdomains and motifs. Domains are colored as follows: N-terminal domain (light gray) with β-hairpin (purple); NiRAN (lime green); interface (green). Subdomains: fingers (dark blue); palm (yellow); thumb (red). Metal ions are shown as spheres, colored lime green (Mg²⁺) and light gray (Zn²⁺). In (B), the RNA duplex is shown with the primer strand in light blue and the template strand in magenta.

Among the structural elements of the polymerase (**Fig. 1**), the thumb subdomain occupies a strategic position linking the RNA-binding regions with catalytic motifs.^2,4,5^ Across viral RdRps, the thumb subdomain mediates large-scale conformational transitions that regulate active site accessibility and catalytic cycling.^6,7^ In contrast, the SARS-CoV-2 polymerase exhibits a structurally distinct, cofactor-dependent organization, suggesting that such regulatory dynamics may be modulated differently or emerge from distributed interactions.^2,5^ This raises the question of whether thumb dynamics in SARS-CoV-2 RdRp play a similar regulatory role, or instead define a distinct mechanism that couples RNA positioning within the channel to catalytic state transitions across the nucleotide addition cycle (NAC; **Fig. 2 top panel**).^8^ Yet how motions within the thumb subdomain contribute to long-range regulation in the apo polymerase remains unresolved. A key open question is therefore how intrinsic apo-state dynamics preconfigure the allosteric networks that govern subsequent RNA engagement and catalysis.

**Figure 2.**
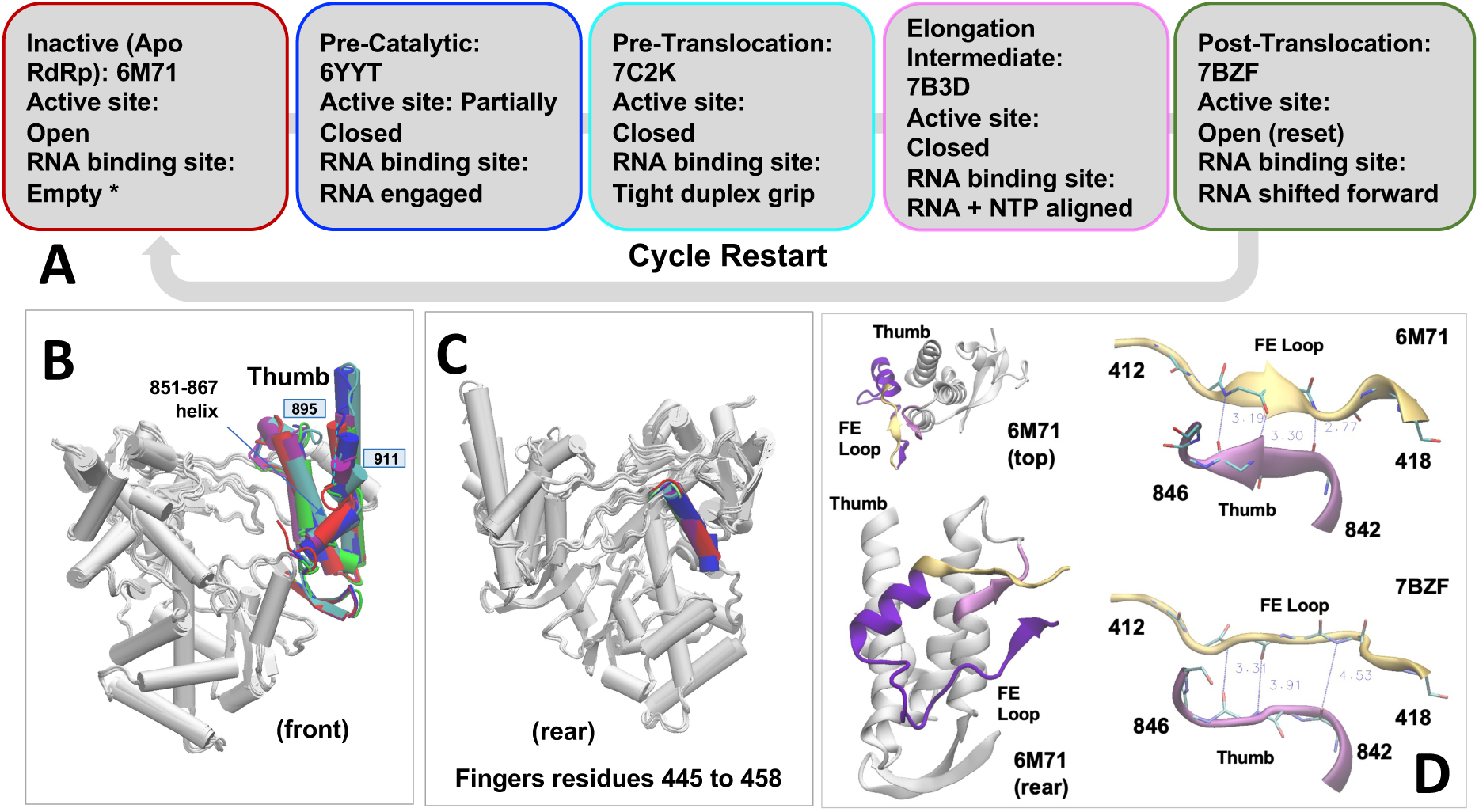
Conformational landscape of the SARS-CoV-2 polymerase across functional states of the nucleotide addition cycle (NAC). (A) Representative structures of the SARS-CoV-2 RNA-dependent RNA polymerase (RdRp) in distinct functional states of the NAC polymerase cycle are shown as boxes colored by functional states, including the apo scaffold (PDB ID: 6M71 in red), RNA-bound pre-catalytic state (PDB ID: 6YYT in blue), catalytic states (PDB IDs: 7C2K in cyan and 7B3D in pink), and post-translocation state (PDB ID: 7BZF in green). These structures illustrate progressive organization of the active site (i.e., the site of catalysis) and engagement of the RNA duplex within the central channel (i.e., the RNA binding site) formed by the fingers, palm, and thumb subdomains. (B)-(D) Structural superposition of representative states highlighting regions of conformational variability observed across the experimental NAC ensemble. Non-overlapping regions are localized to (B) the thumb subdomain, and its interfaces with (C) the fingers (colored by the NAC representatives) and (D) the fingers extension loop (in light pink, yellow and purple, respectively), identifying these regions as the primary sources of structural variation.

Addressing this gap is particularly important because viral RdRps operate through regulatory principles that differ in their structural organization from those of many cellular polymerases. In cellular DNA and RNA polymerases, catalysis and translocation are typically coupled through conformational changes of localized mobile elements, such as fingers-domain motions or O-helix rearrangements.^9,10^ In contrast, viral RdRps are organized as distributed assemblies of conserved structural motifs (A–G) and flexible subdomains (**Fig. 1 Fig. S1**), whose correlated motions contribute to RNA positioning, nucleotide incorporation, and active-site transitions.^1,11–13^ Consistent with computational studies of polymerase systems highlighting distributed allosteric communication networks ^8,14–16^, such coupling is generally described in terms of correlated dynamics spanning multiple structural regions rather than discrete structural switches. Within this framework, protein domains or subdomains such as the thumb can be viewed as components of inter-domain communication pathways linking distal functional regions. In the SARS-CoV-2 RNA-dependent RNA polymerase nsp12, this general principle is preserved but redistributed across distinct structural modules, where thumb-subdomain motions and its coupling with distal domains contribute to long-range allosteric communication and conformational state selection, as described herein.

Although RNA-bound RdRp structures have illuminated catalytic states, they do not reveal how intrinsic motions of the polymerase *predispose* the enzyme toward functional conformations prior to substrate engagement. Ligand binding frequently shifts preexisting conformational populations rather than generating entirely new states, implying that regulatory communication pathways are encoded within the apo enzyme itself. By resolving *how* apo-state fluctuations organize long-range communication across the polymerase, regulatory couplings not apparent from static structures can be uncovered. Revealing how the thumb dynamics influence the enzyme’s functional response to RNA binding while exposing dynamic nodes, may serve as targets for allosteric inhibition of viral replication.

Here, we combine molecular dynamics (MD) simulations with free-energy landscape analysis to resolve how intrinsic motions of the SARS-CoV-2 RdRp organize long-range allosteric communication in the absence of RNA, with particular emphasis on the role of the thumb subdomain. While experimental structures have suggested that cofactor and RNA binding induce subtle thumb-domain rearrangements within a largely preorganized polymerase architecture ^5^, here we define the underlying dynamic and mechanistic basis of this coupling. We show that the apo polymerase samples *preorganized* conformational states in which thumb-subdomain motions dynamically couple catalytic motifs, RNA-binding regions, and distal regulatory elements, including the interface and NiRAN domains, through coordinated collective motions across the conformational free-energy landscape.

Together, these findings provide a dynamic framework for understanding RdRp polymerase nsp12 function and allosteric regulation that complements existing structural models. More broadly, the results identify structurally coupled regions within the coronavirus polymerase that may provide opportunities for conformational and allosteric modulation of viral replication machinery.

## 2. RESULTS and DISCUSSION

### 2.1 Intrinsic Conformational Dynamics of the Apo NSP12 Polymerase Governed by the RdRp-Thumb

We first examined the structure and domain architecture of SARS-CoV-2 nsp12. The polymerase consists of the highly conserved RNA-dependent RNA-polymerase (RdRp) domain (**Fig. 1**), comprising the fingers (blue), palm (yellow), thumb (red) subdomains, as well as an interface domain (lime), and a large N-terminal β-hairpin (grey/purple) extension with a NiRAN domain (dusty-green) with a kinase-like fold^1–5^. The RNA template and product channels are flanked by the RdRp fingers, palm, and thumb subdomains, with the NTP entry site and catalytic motifs A–G positioned at the core (**Fig. S1**). [Herein and throughout the text, the nsp12 domains and RdRp subdomains are referred to collectively as (sub)domains for simplicity.]

Next, we used microsecond-scale molecular dynamics (MD) simulation to characterize the conformational flexibility of apo nsp12 in complex with its essential cofactors nsp8 (2 units) and nsp7 (PDB ID: 6M71)(2) to define the intrinsic dynamic landscape of the polymerase *prior* to RNA engagement. While nsp7 and nsp8 are unique to coronaviruses and closely related nidoviruses, their roles in enhancing processivity, stabilizing RNA, and coordinating replication are functionally analogous to processivity factors and accessory modules employed by other viral and cellular polymerases.^2,17–19^ Structural and evolutionarily closed viral polymerases, such as HCV and poliovirus RdRps, for example, achieve high processivity through intrinsic *enclosure* of the polymerase active site.^6,7^

Integrated analyses of local flexibility, global stability, correlated motions, and low-dimensional conformational sampling (1 *μ*s trajectory) revealed that the apo complex nsp12 populates a structured ensemble of interconverting conformations rather than a single static state. The thumb subdomain emerges as a dynamically responsive “hub-like” region within the conformational ensemble, exhibiting enhanced flexibility while remaining coupled to global structural rearrangements of nsp12 (i.e., the interface domain). Superimposition of experimentally determined structures representative of the nucleotide addition cycle (NAC) ^2,4,19,20^ also highlights the thumb subdomain —together with its interfaces with the fingers subdomain and the fingers extension motif—as the only regions that do not overlap across states (**Fig. 2A-B**), identifying them as the principal loci of structural variation and serving as a reference for the analyses presented below.

#### Local flexibility

Root-mean-square fluctuation (RMSF) analysis of the apo nsp12–nsp7–nsp8.1–nsp8.2 complex reveals heterogeneous flexibility across the polymerase (**Fig. 3A**). Most Cα atoms in nsp12 exhibit limited fluctuations, with the majority remaining well below ∼2 Å and overall values confined to the ∼1–5 Å range (e.g., the N-terminal β-hairpin, NiRAN/interface, palm, and fingers), whereas the thumb exhibits substantially higher mobility, with fluctuations in the ∼3–7 Å range, highlighting it as a dynamically distinct region within the apo ensemble. Notably, this region exhibits intrinsic flexibility, as evidenced by the absence of resolved coordinates across multiple experimental structures, particularly in available apo forms (also highlighted in **Fig. 2A**).^2,4,19,20^

**Figure 3.**
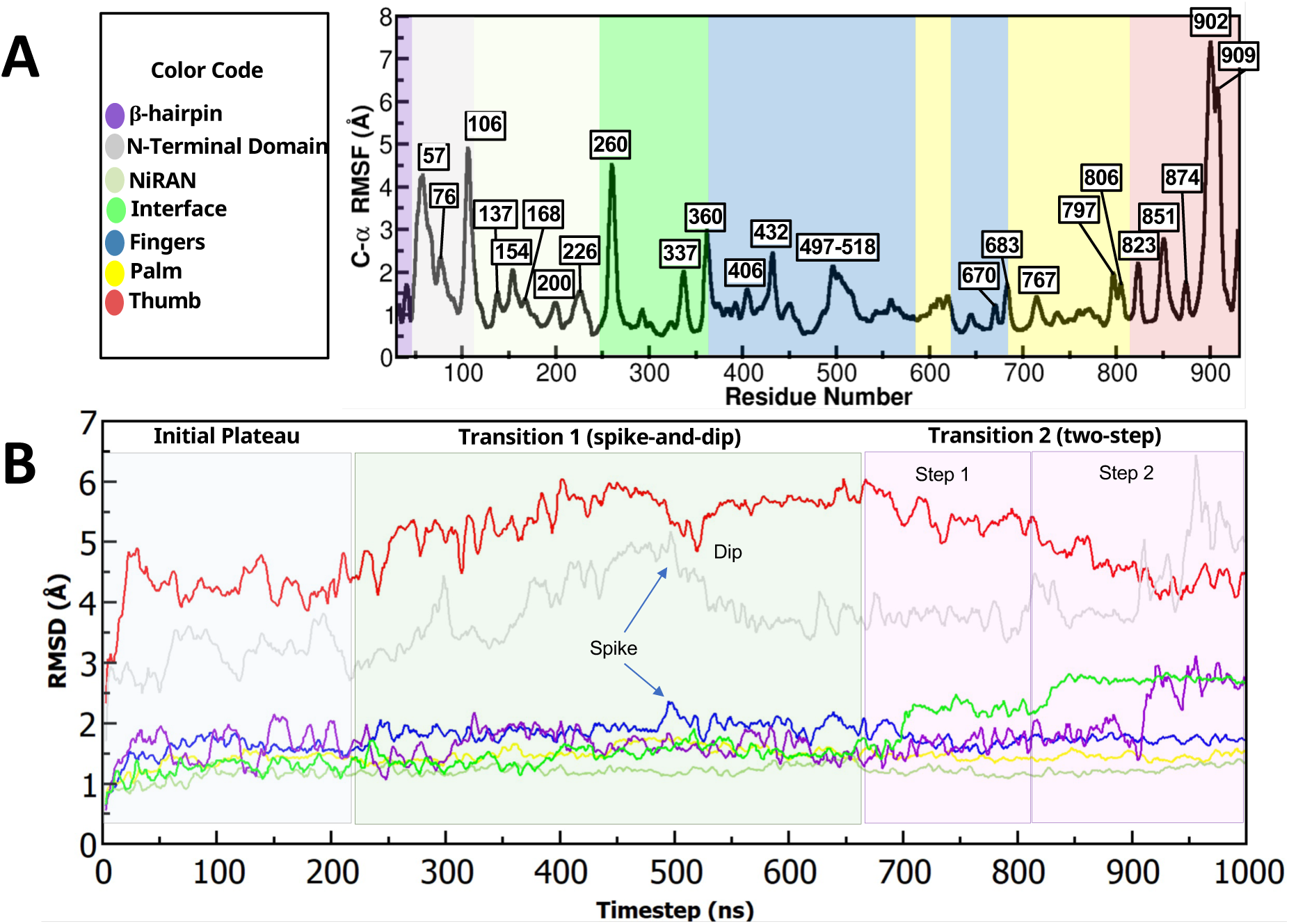
RMSF and RMSD of apo SARS-CoV-2 nsp12. Each nsp12 domain is displayed in the same coloring as in Figure 1. (**A**) Root-Mean-Square-Fluctuation (RMSF) of nsp12 shows average fluctuation of each residue’s alpha carbon (*C*_!_) in angstroms (Å). (**B**) Root-Mean-Squared-Deviation (RMSD) of each nsp12 (sub)domain depicts overall conformational deviation of each (sub)domain from their initial starting conformation over time. One nanosecond is equivalent to one conformation. Running average of 5 was used for all graphs for visualizing smoothed trends of the data.

#### Global Stability & Domain-specific Dynamics

The global backbone (Cα,C,N,O) root-mean-square-deviation (RMSD) of nsp12 is stable across the trajectory (**Fig. S2**), indicating overall structural integrity with a preserved global fold. Notably, a transition in the dynamical behavior of nsp8.1 around 520-530 ns coincides with a subsequent stabilization of the nsp12 trajectory, suggesting a coupled relaxation process at the complex level.^21^ In contrast, the domain-specific analysis reveals a localized behavior (**Fig. 3B**), primarily governed by inter-domain motions within nsp12 that persist even after this global stabilization event. Notably, the local dynamics is primarily governed by the *thumb* region within the RdRp domain (**Fig. 1C**), which exhibits the largest deviations along the trajectory, reflecting its intrinsic flexibility relative to the more structurally stable core of the polymerase. This is followed by the N-terminal domain with *β*-hairpin, with deviations predominantly localized toward the end of the trajectory. Whereas, the remaining domains exhibit more limited fluctuations, with the interface domain representing the most notable exception.

Analysis of the RMSD trace for the RdRp-thumb over the 1-µs trajectory reveals distinct regimes of conformational behavior (**Fig. 3B, red**). **Initial Plateau & RMSD-Transition 1.** An initial plateau (∼0–200 ns), reflecting a relatively stable conformation, is followed by a gradual change in RMSD between ∼200–250 ns into a second, more extended plateau (∼250–650 ns). Within this region of relative stability, a pronounced “dip” at ∼520 ns indicates a brief structural excursion of the thumb subdomain, which correlates with a “spike” in the fingers and the N-terminal domain traces, as well as a notable deviation in the overall palm (**Figs. 3 & S3-S4**), suggesting transient coupling between these (sub)domains within the RdRp polymerase (hereafter denoted as the “**spike-and-dip**” event, ∼520 ns). This event is closely followed by equilibration of the dynamics of the long-fingers/short-palm (**Fig. S3),** and the N-terminal domain with β-hairpin regions, reflecting stabilization into a conformational state that is maintained until the next rearrangement. At the level of global dynamics, this event temporally coincides with the nsp8.1 transition and the ensuing stabilization of nsp12 observed around 520–530 ns (**Fig. S2**). **RMSD-Transition 2.** A discrete **“two-step” dynamical change** occurs at the nsp12 interface domain, with conformational changes detected at approximately 690 ns and 830 ns (**Fig. 3B, green trace**). These events are marked by stepwise increases in the interface-domain RMSD (from ∼1.5 Å to ∼2.2 Å and from ∼2.2 Å to ∼2.5 Å, respectively). Both steps of this RMSD event coincide with coordinated rearrangements in the nsp8 cofactors (**Figs. 3 & S2**), with the most pronounced changes observed in nsp8.2 (step 2). Each rearrangement is also accompanied by a conformational change in the thumb subdomain –the first one starting at ∼670 ns and the second at ∼830 ns– which subsequently adopts a more stable configuration and equilibrates into a distinct, lower-RMSD (from 5.5 to 4.25 Å) conformational state by ∼900 ns (**Fig. 3B**). Concomitantly, stabilization of the thumb during the final 100 ns of simulation is accompanied by increased flexibility of the N-terminal domain with β-hairpin region, whereas the NiRAN domain dynamics remains almost unaffected throughout the trajectory.

These observations indicate that the thumb subdomain samples a set of stable conformational states while undergoing transient, coordinated motions with distal regions of the polymerase, suggesting a *role in interdomain communication*. While most other (sub)domains exhibit comparatively smaller structural deviations, the interface and N-terminal region – the latter showing pronounced fluctuations during RMSD-Transition 2, particularly in the latter part of the trajectory – displays fluctuations over similar time intervals, suggesting participation in shared conformational dynamics. In addition, the interface and β-hairpin domains, although less dynamic overall, show increased fluctuations toward the end of the trajectory, indicating delayed or secondary responses to the dominant thumb motions. Importantly, the magnitude and persistence of the thumb rearrangements distinguish it as the *primary locus* of pronounced intrinsic motion in the apo polymerase. Finally, convergence to stable RMSD plateau further supports equilibration within individual (sub)domains. Notably, the lower final RMSD of the thumb points to a more constrained conformational state relative to other regions, consistent with a functional asymmetry in which this domain stabilizes, while others retain greater flexibility, potentially priming the polymerase for coordinated catalytic activity.

### 2.2 Differential Interdomain Coupling & Conformational Dynamics Reveal “Thumb” Level Coordination

#### Global Domain Coordination and Repositioning

To quantify interdomain rearrangements underlying the observed thumb-driven shifts in RMSD behavior, center-of-mass (COM) distances were calculated between the thumb and the remaining (sub)domains over the course of the simulation (**Fig. 4**). Due to COM distances reporting on **“**global” domain positioning rather than local flexibility, these analyses provide a measure of (sub)domain-level coordination.

**Figure 4.**
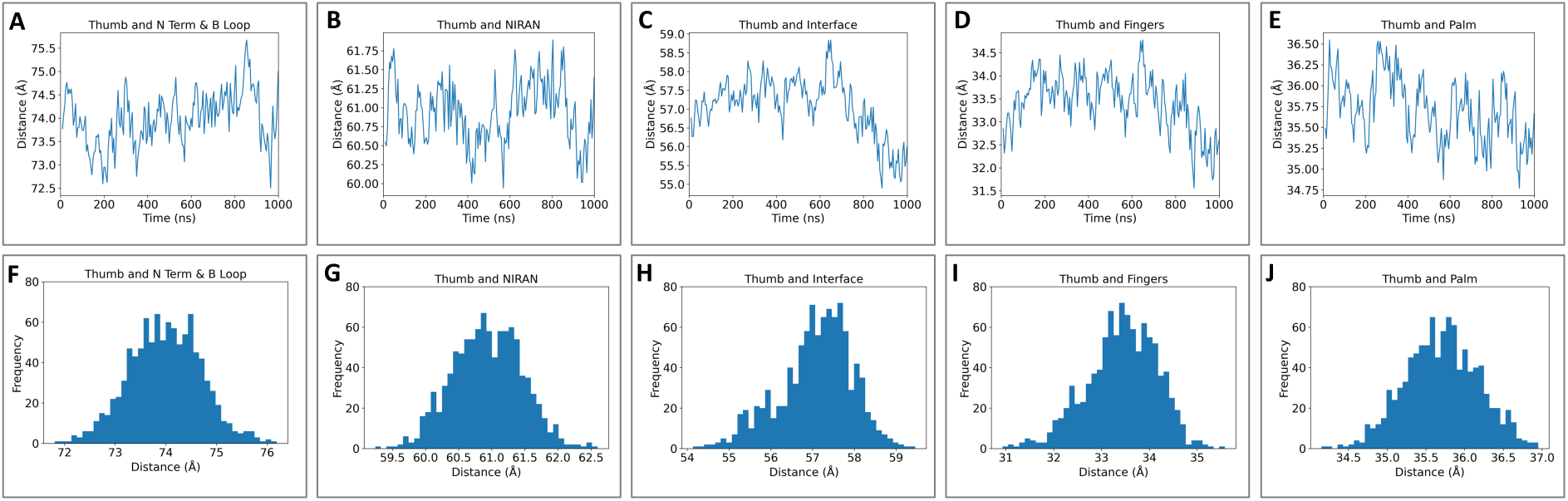
Center-of-mass (COM) distance analysis of inter-domain organization relative to the nsp12-thumb. (A) Time series of COM distances between the thumb subdomain and individual nsp12 (sub)domains (fingers, palm and interface subdomains; NiRAN and N-terminal with *β*-hairpin domains) over the course of the MD trajectory (1 *μ*s), reporting large-scale inter-domain rearrangements. Distances fluctuate on the nanosecond-to-microsecond timescale, with pronounced variability observed for thumb–interface and thumb–fingers separations, while thumb–palm distances remain comparatively constrained. (F)-(J) Probability distributions (histograms) of the corresponding COM distances, quantifying the sampled conformational space. The distributions reveal broad, multimodal profiles for thumb–interface and thumb–fingers distances, consistent with multiple conformational states, whereas narrower distributions for thumb–palm distances indicate a more stable spatial relationship. Overall, comparative representation of COM distance populations highlighting the relative positioning of the thumb with respect to the polymerase (sub)domains. Differences between distributions underscore heterogeneous sampling of inter-domain configurations and suggest preferential states associated with distinct thumb orientations. Together, these analyses define the range and distribution of thumb-centered inter-domain separations sampled in the trajectory.

COM distance analysis (**Fig. 4)** reveals that the thumb subdomain maintains structured but differential coupling with the remaining nsp12 (sub)domains. Specifically, the distance between the thumb and NiRAN domains shows a relatively narrow, roughly unimodal distribution (around ∼60–62.5 Å) and remains stable, with no major state switching. This suggests that the thumb and NiRAN domains move in a strongly coupled, coordinated manner both structurally and dynamically. Such stable inter-domain coupling is consistent with structural studies showing the NiRAN–thumb interface as a rigid core in the RdRp complex,^2^ proposing limited relative motion in the apo form. Next, the distance between the thumb and interface exhibits a broader, slightly skewed distribution that gradually decreases over time. The COM profile for this pair shows a clear time-dependent shift rather than stable fluctuations, indicating that their relative positioning changes during the simulation. A drift after ∼700 ns coincides with the first event in the two-step rearrangement observed in RMSD analyses, showing a pronounced temporal change in the COM distance and indicating that the thumb and interface move together during this dynamic change (**Fig. 3B**, interface RMSD trace). This pattern of slow domain rearrangement aligns with dynamic behavior observed in MD and structural studies of CoV polymerases undergoing conformational transitions prior to catalysis^2^; consistent with coordinated motion between functional domains. Next, after a relatively fast increase around ∼200 ns, the distance between the thumb and fingers domains stabilizes, compatible with the timing of the second RMSD plateau with spike-and-dip event (**Fig. 3B**, fingers RMSD trace), before gradually decreasing again after ∼700 ns. This behavior indicates progressive convergence of these domains, with the fingers participating in adaptive motions alongside the thumb, potentially contributing to allosteric propagation. Comparable convergence of thumb and fingers domains upon engagement with substrate has been observed in active RdRp structures and related simulations.^5^ In contrast, the COM distance between the thumb and palm domains exhibits a relatively broad distribution (∼34–37 Å) with no evidence of a sustained long-timescale drift, but rather higher variability, consistent with a more flexible, loosely coupled relationship in the apo state. Such variability accords with structural and simulation studies of CoV RdRp in its pre-engagement form, where the polymerase adopts a relaxed, open conformation prior to RNA/NTP binding, and the thumb is not yet engaged as a stabilizing clamp on the palm.^2^ Lastly, COM distance analysis between the thumb and the N-terminal domain with β-hairpin shows substantially larger separations (∼72–76 Å) compared to the other domain pairs, with moderate fluctuations and a unimodal distribution centered near ∼74 Å. In contrast to the coordinated and directional behavior observed for the other domains, the distance between the thumb and the N-terminal/β-hairpin domains is not monotonic; indicating the absence of concerted global repositioning and suggesting that the N-terminal/β-hairpin domain remains spatially distal and exhibits only limited coordination with thumb-driven motions. The relatively stable average distance, despite thermal fluctuations, indicates that this domain maintains a consistent global orientation while not participating in the coordinated conformational rearrangements that modulate RNA-channel geometry. This behavior is consistent with structural studies showing increased flexibility of the N-terminal/β-hairpin region relative to the polymerase core.^19^

COM distances relative to the full RdRp were also monitored (**Fig. S5**). Specifically, the interface domain exhibits a time-dependent shift in its COM distance relative to the polymerase (bimodal distribution, strong signal), consistent with progressive repositioning during the observed dynamic changes. In contrast, whereas NiRAN remains rigidly anchored (narrow unimodal distribution, minimal shift), the N-terminal domain fluctuates without directional coupling (broader, yet stable, distribution).

These observations suggest that the RdRp *thumb acts as a dynamic integrator,* coordinating both stable and adaptive interdomain interactions across the polymerase. Together, the results reveal a hierarchy of interdomain coupling, in which thumb motions strongly coordinate proximal elements such as the fingers, moderately influence adjacent regions, and have limited impact on distal domains. The observed dynamics are dominated by relative interdomain rearrangements rather than large-scale compaction, highlighting that structural variability arises primarily from coordinated repositioning of individual domains. This spatially organized pattern of coupling supports a model in which the thumb orchestrates global polymerase rearrangements.

#### Dynamic Cross Correlation: Strength and Pattern of Inter-Domain Coupling

To further dissect coordinated motions across the polymerase, we next examined dynamic cross-correlation (DCC). The global DCC map reveals distinct patterns of correlated and anticorrelated motions across nsp12 (sub)domains (**Fig. 5A**). The thumb subdomain behaves as a coherent unit within the RdRp polymerase core, showing positive correlations with the fingers and portions of the palm (long fingers and short palm regions), and anticorrelations with the N-terminal/β-hairpin region. The fingers and palm subdomains display strong internal correlations, consistent with coordinated motions within the polymerase core. The interface domain exhibits a heterogeneous correlation pattern, with regions showing positive correlation with the thumb and fingers, and other regions displaying anticorrelated behavior relative to the N-terminal/β-hairpin and parts of the palm. The N-terminal and NiRAN regions show internally correlated motions, with weaker and more variable coupling to the core domains.

**Figure 5.**
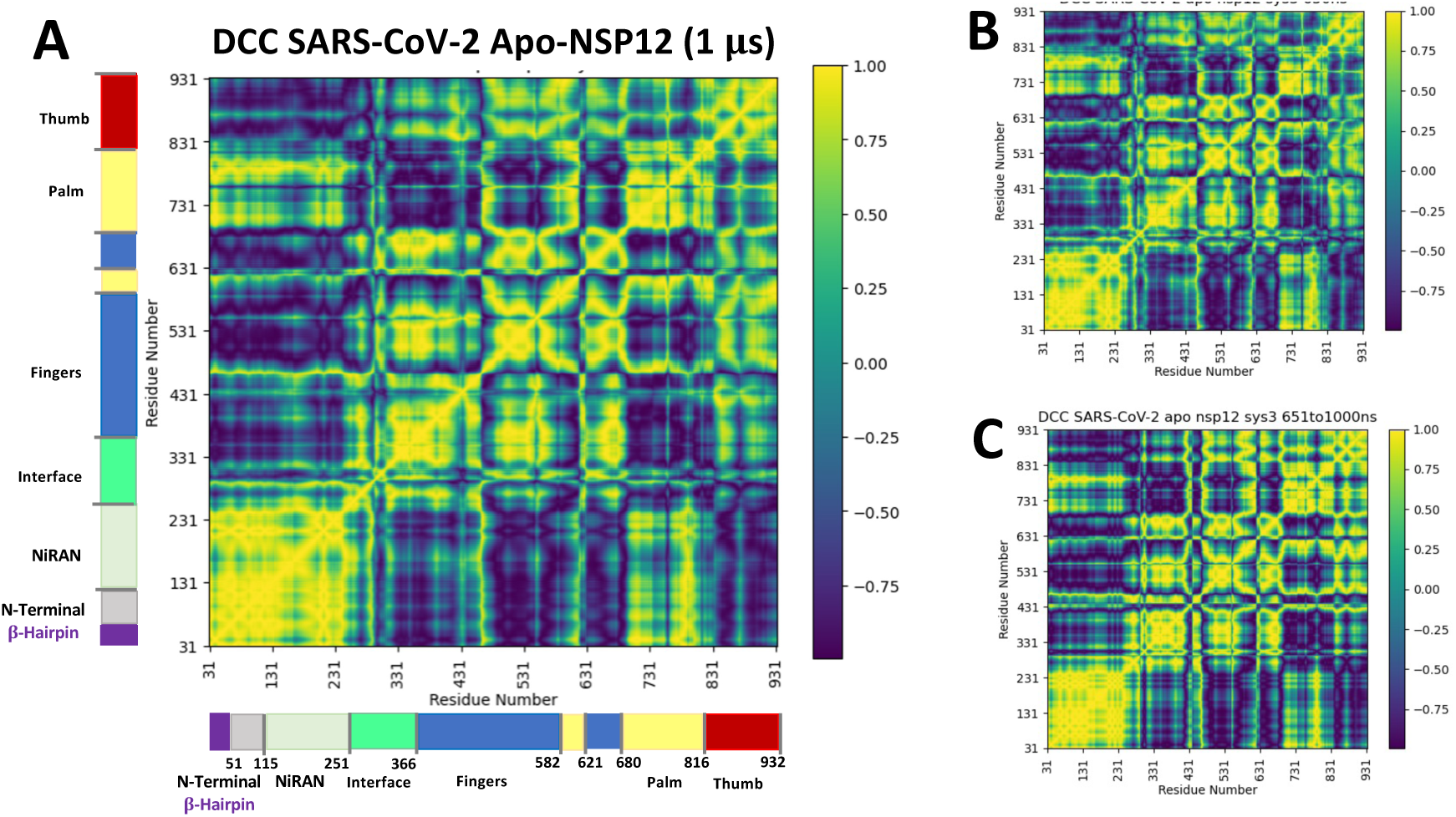
Dynamical cross-correlation analysis reveals transient rewiring of domain communication. (A) Global dynamical cross-correlation (DCC) map averaged over the full trajectory (1 *μ*s), showing persistent correlated (yellow) and anti-correlated (purple) motions across the polymerase. (Sub)Domain boundaries (N-terminal β-hairpin, NiRAN and interface domains; fingers, palm and thumb subdomains) are indicated with coloring consistent with Fig. 1. While the global map captures overall (sub)domain organization and long-range coupling, it reflects a time-averaged view of the dynamics. (B) Time-resolved DCC map computed over a sliding window (1–650 ns), revealing heterogeneous and dynamically evolving coupling. In contrast to the global map, correlations are fragmented and reorganize over time, with transient interactions particularly involving the interface and thumb regions and variable coupling with fingers and palm. (C) Time-resolved DCC computed over a sliding window (651–1000 ns), emphasizing recurrent correlation features, identifying reproducible patterns of inter-domain coupling. These data indicate that the system samples discrete dynamical regimes distinguished by specific interface–thumb and polymerase core interactions. Together, the time-resolved analyses demonstrate that the correlation network is dynamically reconfigured rather than static, uncovering transitions that are not apparent in the global map.

In contrast, time-resolved DCC analysis reveals a transition in thumb–interface coupling. In the early portion of the trajectory (∼first 650 ns), the thumb and interface regions are predominantly positively correlated, whereas in the later portion (∼ last 350 ns) they become anticorrelated (**Fig. 5B–C**). Intra-domain correlations within the thumb, as well as its coupling to the fingers and palm subdomains, remain largely unchanged across both time windows. This temporal transition is consistent with changes observed in COM distance (**Fig. 4C–H**) and RMSD analyses affecting the polymerase stability both at the subdomain (**Fig. 3B**) and the global (**Fig. S2**, nsp12 RMSD trace) levels.

Overall, these patterns indicate that thumb motions are preferentially coupled to proximal subdomains (i.e., the fingers and palm), while distal regions exhibit weaker or opposing correlations, supporting a spatially organized hierarchy of interdomain coupling.^2^ These correlation patterns are consistent with domain-level reorganization observed in COM and RMSD analyses, linking coordinated motions to underlying changes in dynamical regimes.^19^

#### Distinct Coupled states and Structural Rearrangements

**Principal component analysis (PCA) of the MD backbone ensemble**^22^ shows the distribution of frames projected onto the first two principal components (PC1 and PC2) for the nsp12 protein, the RdRp domain, and the thumb-subdomain datasets, respectively (**Fig. 6**). In all three projections, the sampled configurations occupy multiple visually distinguishable clusters in PC1–PC2 space. The cumulative variance plot indicates that the first three principal components account for approximately ∼50% of the total variance for NSP12 (PC1–PC3), ∼55% for RdRp, and ∼65–70% for the thumb dataset, with a progressively more concentrated trend upon domain reduction (**Table S1**). This is in line with previous observations: (i) RMSD trajectories show plateau regions in selected components, including a two-step RMSD pattern in the thumb region (**Fig. 3B**); (ii) RMSD-derived DCC matrices indicate correlated residue fluctuations across structural domains dominated by thumb motions (**Fig. 5**); and (iii) COM distance distributions between key regions show multimodal behavior, including a bimodal distribution for interface–thumb distances (**Fig. 4**). Furthermore, PCA of NiRAN, interface, and combined palm/fingers shows the interface contributes a large variance fraction (∼66%) (**Fig. S6**), comparable to what observed with the thumb dubdomain (**Fig. 7**). This is accompanied by the RMSD two-step behavior and bimodal COM distributions between interface and thumb (**Figs. 3B, 4C and 4D**). PCA subspace projections of the interface show separation consistent with these distributions.

**Figure 6.**
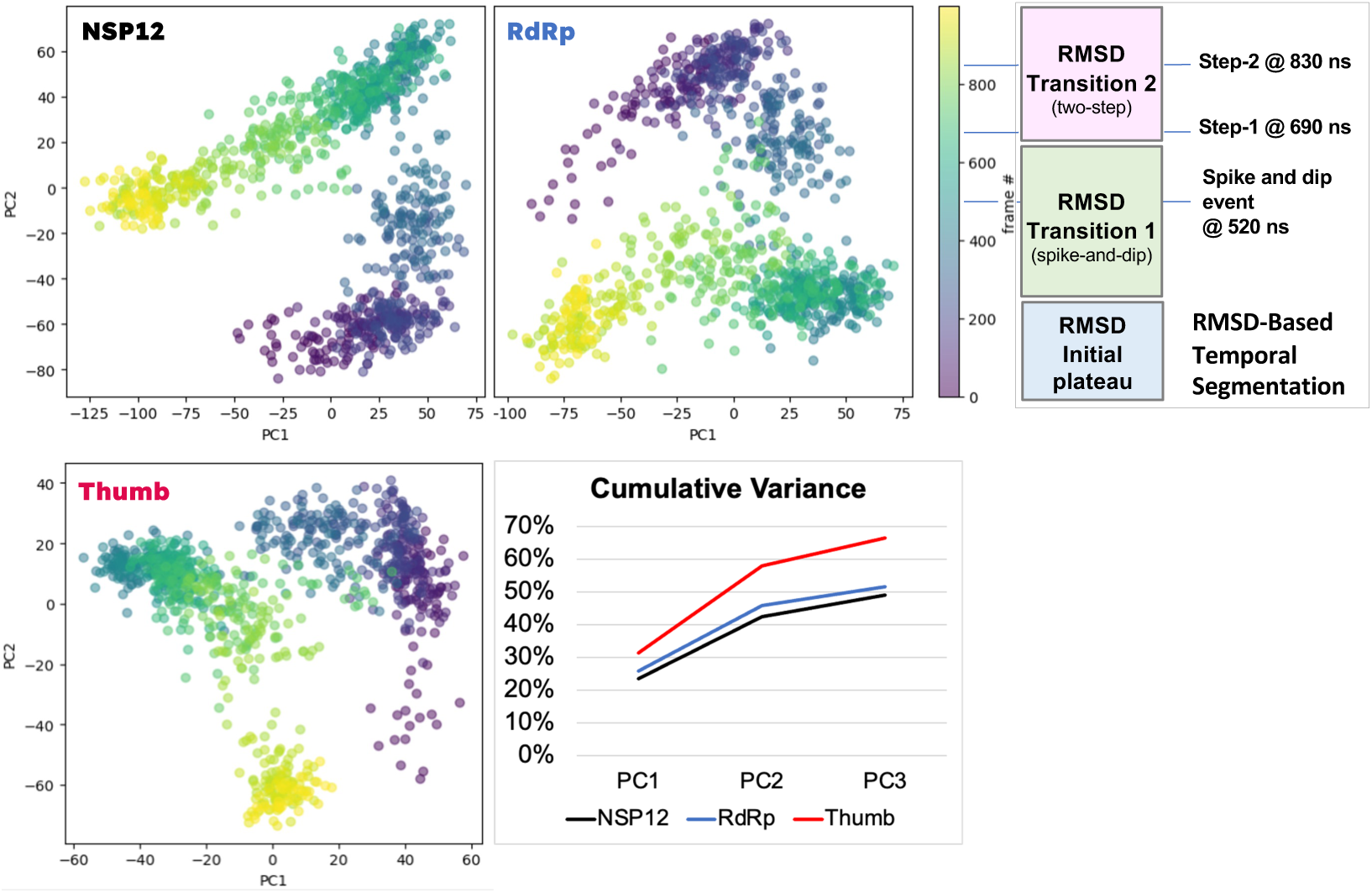
Principle Components Analysis (PCA) and Cumulative Variance of apo nsp12, RNA-Dependent RNA Polymerase (RdRp), and the Thumb Subdomain. The PCA graphs, labeled for each structure analyzed with color scheme from Figure 1, shows the transformed data using two principal components for each. These uncorrelated points are colored by frame number using the gradient at the right of the graphs (each frame corresponds to 1 ns, for a total of 1000 frames or 1 *μ*s). The fourth plot at the bottom right is the cumulative variance for each analyzed structure overlaid to demonstrate the enrichment as the structure analyzed becomes specific. RMSD-based temporal segmentation is also shown for comparison with PCA projections (top right). Schematic view of major conformational regimes identified from the RMSD trajectory, included to provide temporal context for the PCA analysis in which frames are colored by simulation time. The system initially resides in a stable initial plateau, followed by Transition 1, characterized by a spike-and-dip event at ∼520 ns. A subsequent two-step transition (Transition 2) occurs, with changes at ∼690 ns (Step 1) and ∼830 ns (Step 2). The vertical axis indicates simulation time (frame number). This scheme enables direct mapping of RMSD-defined transitions onto the time-colored PCA projections, facilitating interpretation of how large-scale structural rearrangements are distributed in principal component space.

**Figure 7.**
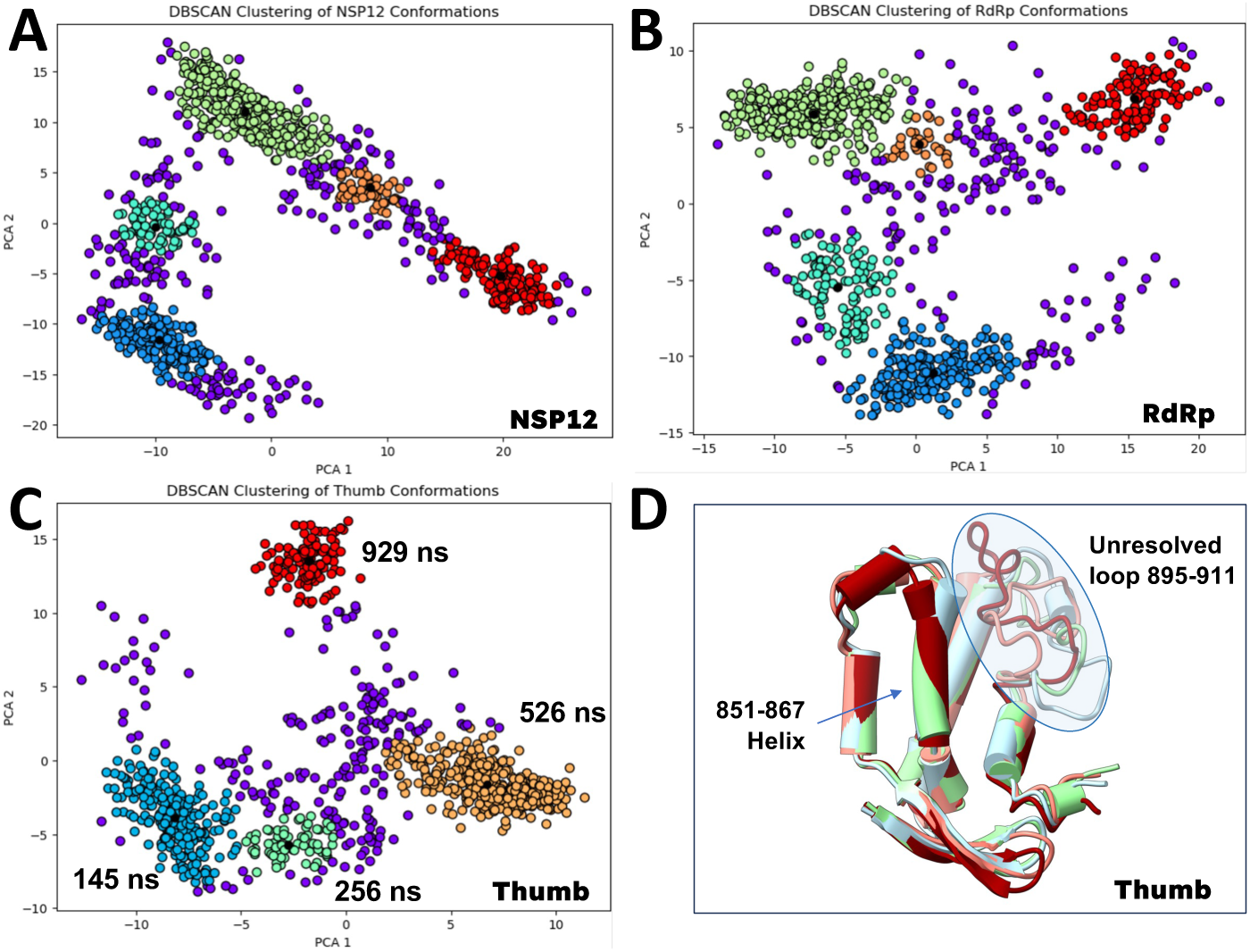
Density-Based Spatial Clustering of Applications with Noise (DBSCAN) of Apo NSP12. Apo nsp12 RNA-dependent RNA polymerase (RdRp) and the isolated thumb subdomain are shown (A to C, respectively). DBSCAN applied to each system identifies clusters of structurally similar conformations. Purple points denote noise/outliers not assigned to any cluster, while black markers indicate cluster centroids (representative structures). (D) Superposition of the thumb subdomain of representative clusters from (C). The 895-911 loop (also in Fig. 2A) unresolved across the experimentally available nsp12 structures is highlighted (oval). A helix exhibiting pronounced conformational flexibility (residues 851-867, also in Fig. 2B) is also shown.

**Clustering by DBSCAN**^23^ (**Fig. 7**) of PCA coordinates identifies discrete clusters across full-length nsp12, RdRp, and thumb systems. The full-length and RdRp datasets populate multiple distributed clusters in PC space, while the thumb dataset shows more compact cluster separation, as corroborated by superimposed representative frames (**Fig. 7D**). **Clustering by K-means**^24^ (**Fig. S8**) yields the same partitioning. Together, results from PCA and clustering indicate that nsp12 dynamics are organized into a finite set of conformational states, with increasing coherence and dimensional reduction upon focusing on the thumb subdomain. This suggests that the thumb captures a dominant component of the conformational state architecture of the polymerase. Together, PCA-based clustering (DBSCAN and k-means), RMSD profiles, RMSD-DCC patterns, and COM distributions provide consistent partitioning of conformational space across nsp12, RdRp, and thumb systems.

#### Per-residue contributions mapped onto conserved motifs and loop regions

Normalized amplitudes of residue-wise contributions to the first (PC1, blue) and second (PC2, orange) principal components derived from PCA of the trajectory (*C*_!_ atoms) identify the regions that drive the principal dynamical modes of the system (**Fig. S7**). Globally, PC1 exhibits distributed contributions across the sequence, with prominent peaks in the N-terminal/NiRAN/interface regions and a secondary increase toward the thumb subdomain, consistent with large-scale coordinated motions. In contrast, PC2 shows more localized contributions, primarily confined to the N-terminal region, with minimal involvement of the polymerase core. Conversely, residues within the central RdRp domain (and related subdomains, **Fig. 1**) display comparatively low amplitudes in both components, indicating limited participation in the dominant collective motions.

#### Residue-resolved PCA loadings were also mapped onto conserved motifs and annotated loop regions of nsp12^5^

Motif G (499–511), Motif F (544–560), Motif A (612–626), Motif B (678–710), Motif C (753–767), Motif D (771–796), and Motif E (810–820) all exhibited comparatively low normalized amplitudes in both PC1 and PC2 across their respective sequence ranges. These regions correspond to conserved catalytic and RNA-binding motifs that define the structurally ordered RdRp core described in cryo-EM and structural studies of coronavirus polymerases.^2,4,19,20^ In contrast, the finger extension loop (401–447) displayed elevated amplitudes in PC1 together with measurable contributions in PC2, whereas the highly flexible, experimentally unresolved thumb region (∼906–929) showed increased amplitudes predominantly in PC1. Both regions correspond to peripheral structural elements implicated in inter-domain and cofactor-associated contacts,^4,12^ and were highlighted in our experimental (NAC representatives) and MD (clustering) ensembles analysis (**Figs. 2B-2D**, and **7D**).

Across all annotated regions, PC1 contributions were broader and higher in magnitude, whereas PC2 contributions remained comparatively localized and reduced. Together, these residue-wise PCA loading distributions, derived directly from the MD trajectory, define the relative contributions of conserved motifs and loop regions to the dominant collective motions of nsp12.

### 2.3 Inter-domain Rearrangements Retain a NAC-Polymerase Relevant Conformational Ensemble

#### Radius of Gyration (Rg) of the RNA-binding Channel Underlies NAC Relevant RdRp Sampling

A key conceptual distinction highlighted in the NAC framework (**Fig. 2A**) is the difference between the active site and the RNA-binding site. The active site is a localized catalytic pocket, defined by conserved sequence motifs that dynamically open and close to enable nucleotide incorporation and metal coordination. In contrast, the RNA-binding site is an extended structural feature, consisting of a continuous, positively charged channel that spans the polymerase and remains engaged with RNA throughout elongation.^5^ While the active site governs the chemical step of bond formation, the RNA-binding channel controls substrate positioning, duplex stability, and translocation. Thus, efficient RNA synthesis depends on tight coupling between these two layers: local catalytic rearrangements occur within the active site, while global RNA handling and movement are orchestrated by the surrounding channel. This distinction is illustrated schematically in **Figure 2**, where the active site is depicted as a dynamic, state-dependent element, whereas the RNA-binding path persists across all RNA-bound states but varies in its mode of engagement. To assess whether the interdomain motions observed in our MD study affect the structural integrity of the functional polymerase site, the radius of gyration (Rg) was calculated for residues defining the RNA-binding channel as defined in Ref. ^5^ thereby tracking the spaxal distribuxon of these residues throughout the trajectory (**Fig. 5A**). The Rg trajectory was then compared against Rg values calculated for representative experimental structures from the SARS-CoV-2 NAC cycle (**Fig. 2**), indicating that the polymerase is dynamically regulated without loss of catalyxc competence.

#### Rg of Experimentally Determined NAC States

The apo state (PDB-ID 6M71)^2^ in the original experimental structure (**Fig. 5**) exhibits the second lowest Rg (∼15.5 Å, **Table S2**), indicating a relatively compact or collapsed channel in the absence of RNA. Upon RNA binding and during the pre-translocation (16.04 Å) and post-translocation (15.94 Å) events, Rg increases, reflecting structural expansion or rearrangement of the channel to accommodate the RNA duplex (PDB-IDs 7C2K and 7BZF, respectively).^4^ Notably, the Remdesivir-stalled state (7BV2) exhibits a similar Rg (∼15.84 Å), consistent with the pre-catalytic 6YYT^19^ (∼15.82 Å) and elongating (7B3D)^20^ (∼15.76 Å) polymerases, suggesting that these functional states share a comparable global compactness. The backtracked polymerase (7KRN)^25^ also exhibits similar Rg (∼15.78 Å), consistent with preservation of overall global compactness despite local rearrangements associated with RNA backtracking. Finally, the capping intermediate state (7THM)^26^ returns to a more compact channel (∼15.40 Å, lowest Rg), possibly reflecting reorganization after RNA processing. Overall, this experimental trend (**Table S2**) supports the view that the RNA-binding channel is dynamic, expanding to accommodate RNA and related intermediates, rather than simply closing xghtly around the substrate. This expansion is essenxal for the polymerase to process RNA efficiently through different stages of the catalyxc cycle.

#### Rg of Apo NSP12 Trajectory

In contrast to the domain-level rearrangements revealed by RMSD, DCC and COM analyses (**Figs. 3-5**), the Rg of the RNA binding site **(Fig. 8)** remains fairly stable throughout the simulation, with moderate fluctuations but no sustained drift. This indicates that, during global conformational changes **(Figs. 4-6)**, the site retains its overall compactness while dynamically fluctuating within biologically relevant ranges consistent with those observed for experimental NAC states (**Fig. 8A-B**). Because Rg reflects local structural organization at the RNA-site level, these results suggest that the observed thumb-dominated global motions in the polymerase modulate the channel geometry without disrupting the structural framework required for catalysis. The observed short-timescale fluctuations likely reflect dynamic breathing of the channel, consistent with functional flexibility rather than structural instability.^5,6^ Furthermore, a temporal shift in the average Rg is observed concomitantly with the onset of the two-step RMSD transition event (**Fig. 3B**), along with the decrease in the thumb–interface COM distance (**Fig. 4C,H**). These changes collectively indicate a structural compaction event supported by correlated thumb–interface motions (**Fig. 5B**, time-resolved DCC).^5^ [Across the 1 μs simulation, the average Rg is 15.73 Å, as seen in **Fig. 8A**. Time-resolved analysis shows a decrease in Rg from 15.77 Å to 15.65 Å, corresponding to the system before and after the two-step interface event (∼650 ns), respectively, indicating a gradual global compaction.]

**Figure 8.**
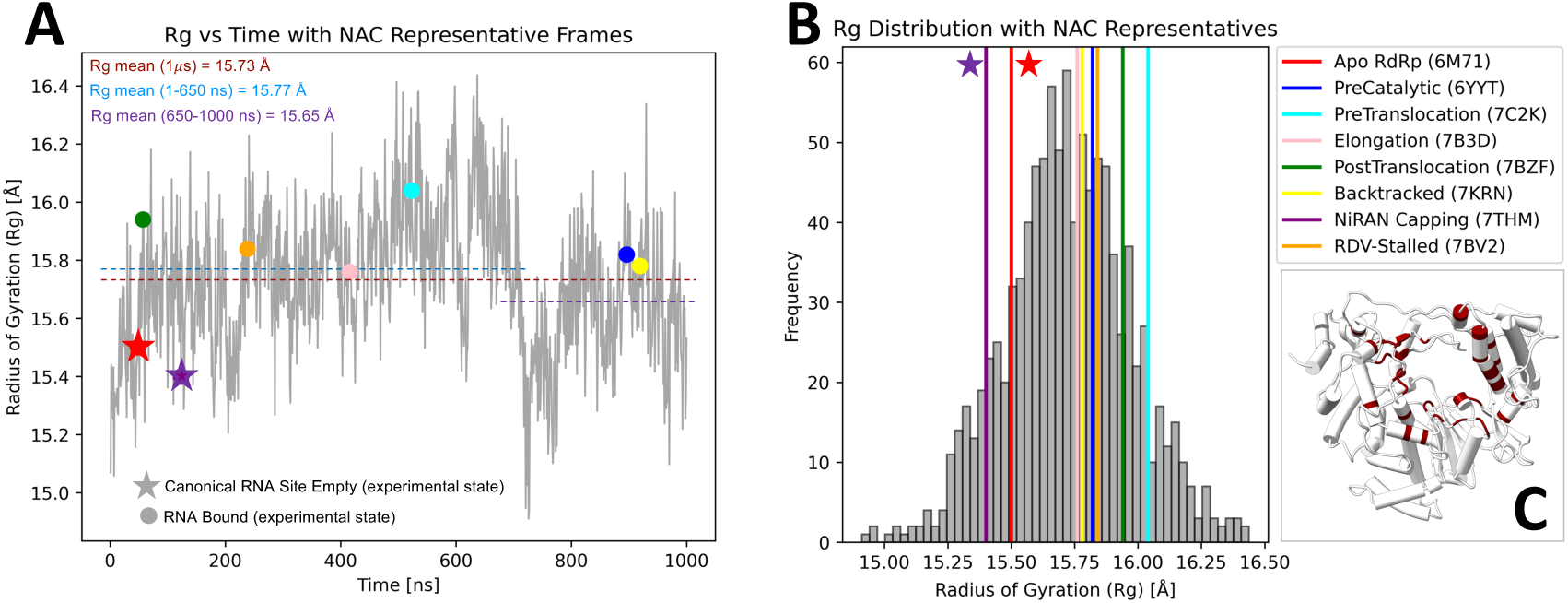
Radius of Gyration (Rg) of RNA Binding Site of the Apo NSP12-RdRp Ensemble and NAC Representative Conformations. (A) Time series of calculated radius of gyration (Rg) values across the MD trajectory for residues in contact with RNA. Colored points indicate frames closest to each NAC representative conformation identified from the experimental ensemble. NAC representatives are color-coded consistently across all panels and correspond to those shown in Figure 2. Additional experimentally derived conformations are overlaid for comparison. (B) Distribution of Rg values calculated over the full trajectory shown as a histogram. Vertical colored lines indicate the Rg values corresponding to NAC representative conformations, while additional markers denote experimentally characterized conformational states (see Legend). (C) RdRp polymerase structure of a randomly picked representative frame shown in white. Residues in contact with RNA), used to calculate Rg values, are highlighted in maroon. The panels illustrate the biological relevance and the extent of conformational sampling captured within the simulation ensemble.

Collectively, these analyses converge on a consistent picture of coordinated, functionally relevant dynamics. Rg variations remain small and within the range of experimentally observed conformations (both within NAC states and additional experimental complexes), while their temporal alignment with RMSD changes events supports structured (sub)domain rearrangements rather than stochastic drift. DCC analysis reveals a coherent correlation network, with thumb–fingers/palm coupling and N-terminal anticorrelation, indicative of coordinated motion. This is further supported by COM trajectories showing a hinge-like displacement of the interface synchronized with the thumb, leading to progressive interdomain closure.^5,6,21^

Notably, apo-like (empty) structures consistently populate lower Rg values, whereas RNA-bound states occupy higher Rg regions across functional states (**Fig. 8 A-B**, “stars” versus “dots”), indicating a robust Rg-dependent separation of experimentally resolved conformations. A consistent shift is also observed in the thumb–interface COM distance (**Table S2**), which further discriminates apo and RNA-bound states, supporting its role as an additional orthogonal collective variable. Together, these features motivate the use of Rg and thumb–interface COM distance for classification of MD-derived ensembles.

### Rg–COM State-space Classification of Apo NSP12 Suggests “Preorganized Thumb States” for Antiviral Targeting

#### Time resolved conformational sampling, NSP12 populations & NAC states

Projection of the MD trajectory onto the Rg–COM coordinate space revealed a structured conformational distribution with an elongated topology reflecting strong coupling between RNA-binding site compactness and thumb–interface separation (**Fig. 9A**). Most frames cluster within an intermediate region centered near Rg values of ∼15.5–16.0 Å and COM distances of ∼56.5–58.5 Å, while lower-populated regions extended toward both more compact and more expanded conformations. Time-resolved coloring show that early, intermediate, and late trajectory frames occupy overlapping but gradually displaced regions of state space (with temporal separation comparable to that seen for RMSD), suggesting progressive sampling across neighboring conformational regions during the simulation. The ensemble does not distribute uniformly across the projection, but instead forms locally dense populations separated by lower-density regions. Given the choice of the collective variables, this dynamic behavior reflects a low-dimensional conformational manifold that gates access across the polymerase through thumb–interface coupling.

**Figure 9.**
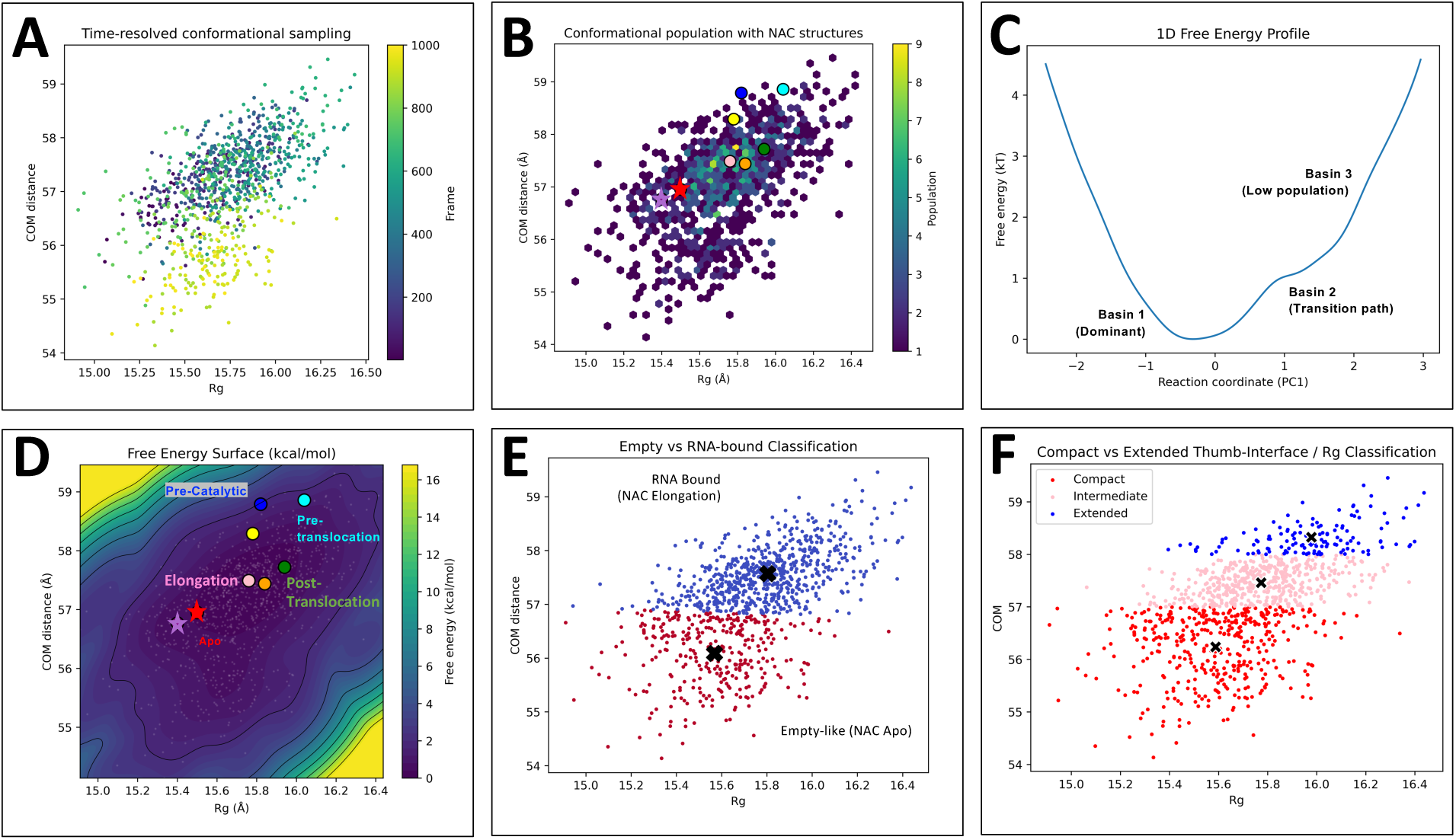
Analysis & Classificadon of Apo NSP12 Conformadonal States Using Rg–COM State Space Analysis. In all panels, the conformational ensemble is projected onto radius of gyration (Rg) and center-of-mass (COM) distance coordinates. While Rg represents the compactness of the RNA-binding site, COM corresponds to the thumb–interface domain separation. (A) Scatter plot of Rg versus COM. Individual points represent trajectory frames and are colored according to simulation time progression. The distribution spans a continuous but nonuniform region of conformational space, with sampling concentrated between Rg ≈ 15.4–16.1 Å and COM distances ≈ 56.5–58.5 Å. Temporal coloring indicates progressive exploration of neighboring regions within the sampled conformational landscape over the course of the trajectory. (B) Hexagonal occupancy map of the conformational ensemble. Color intensity reflects the number of trajectory frames populating each region of conformational space, highlighting dominant conformational basins sampled during the MD trajectory. (C) One-dimensional free energy profiles reconstructed from KDE-derived probability distributions for COM distance and Rg. Free energies are reported relative to the global minimum and reveal a dominant compact conformational basin (minimum) together with a lower-population extended state connected through an intermediate transition region (shoulder). (D) Free energy surface (FES) projected onto Rg and COM distance coordinates. Colors represent relative free energy, with low-energy regions corresponding to preferentially sampled conformational states. Contour lines indicate the underlying energy landscape, while white points denote sampled MD conformations. Experimental structures and NAC representatives are indicated as dots or stars, corresponding to RNA-bound or empty (apo-like) conformations, respectively, while additional markers denote experimentally characterized conformational states (see Fig. 8 legend). (E) Projection of the conformational ensemble onto the Rg–COM distance, colored according to two clusters obtained using k-means (k = 2). Red and blue points represent two major conformational populations, designated here as “empty-like” and “RNA-bound-like,” respectively. Black “X” markers indicate the corresponding cluster centroids. Cluster assignments were determined by mapping each centroid to the nearest experimentally derived NAC representative structure in the Rg–COM space. The centroids were found to correspond most closely to the Apo and Elongation experimental states, respectively, with centroid values being extremely close to those observed for these NAC reference structures. (F) Projection of the conformational ensemble onto the Rg–COM distance, with structures classified into Compact, Intermediate, and Extended conformational populations shown in red, pink, and blue, respectively. Classification was performed using threshold-based assignment along the COM coordinate (57.0 & 58 Å, see Table S2 for comparison with NAC structures). Black “X” markers indicate the mean Rg–COM coordinates of each conformational population, representing the average position of the Compact, Intermediate, and Extended states within the sampled ensemble.

Furthermore, the hexagonal occupancy map (**Fig. 9B**) revealed a non-uniform distribution of conformational sampling, with high-intensity regions indicating dominant conformational basins frequently occupied during the MD trajectory, while lower-intensity regions corresponded to less populated transitional conformations. Experimental structures, shown as individual dots or stars, map predominantly onto the most populated conformational basins of the simulated ensemble.

#### Rg–COM Free Energy Landscape

**1D Free-energy Plot (Rg–COM)**: The 1D free-energy surface projected onto Rg and COM distance (**Fig. 9C**) reveals a dominant low-energy basin corresponding to compact NAC conformations (Basin 1, PC1 minimum −0.5 to 0), with a continuous free-energy gradient toward higher Rg and COM values indicative of progressively expanded NAC states. The landscape lacks a sharp separation between states, instead suggesting a shallow metastable shoulder (Basin 2, PC1 >0.8) and a continuum of conformational rearrangements (Basin 3, high tail).

#### 2D Free-energy Plot (Rg–COM)

The 2D free energy surface (**Fig. 9D**) projected onto Rg and COM distance revealed a broad dominant low-energy basin centered at intermediate Rg (∼15.6–15.9 Å) and COM distances (∼56.5–58.0 Å), corresponding to the most stable conformational ensemble sampled during the simulation. The elongated shape of the basin suggests a continuum of structurally related conformations rather than discrete state transitions. Higher free-energy regions extended toward both more compact and more expanded conformations, indicating reduced sampling of these states. Experimental structures, represented as individual dots or stars, mapped predominantly within the major low-energy basin, supporting agreement between the simulated ensemble and experimentally resolved NAC-related conformations. Specifically, NAC analysis groups structurally characterized RdRp conformations into three broader dynamical classes: (i) a pre-engaged/capping-associated state including apo and NiRAN-capping conformations; (ii) an elongation-competent forward-translocated state encompassing canonical elongation, post-translocation, and remdesivir-stalled complexes; and (iii) a catalytically paused state including precatalytic, pre-translocation, and backtracked conformations. While both states (ii) and (iii) constitute RNA bound polymerase complexes, state (i) feature empty RNA sites.

#### Rg–COM State Classification

The relevance of the Rg–COM state space was further validated by the localization of experimentally resolved NAC conformations, including both apo/empty and RNA-bound structures, within the major populated basins. Accordingly, trajectory frames were subsequently classified based on their Rg and COM values to distinguish compact, intermediate, and extended conformational states and relate them to experimentally observed empty versus RNA-bound conformations, as described below.

#### Empty versus RNA-Bound Classification

K-means^24^ clustering (k=2) in the Rg–COM conformational space identified two major ensembles separated by distinct compactness and (sub)domain-opening characteristics (**Fig. 9D**). The resulting cluster centers (Rg, COM: 15.57 Å, 56.09 Å; and 15.80 Å, 57.57 Å), located near the apo and the elongation states, respectively (**Table S2**), captured the dominant partitioning between a compact closed-like ensemble and a more expanded RNA-associated ensemble. Comparison with experimentally derived NAC reference structures showed that the compact cluster aligned closely with the apo/empty state, whereas the expanded ensemble encompassed conformations associated with RNA-bound functional states.

#### Compact versus Extended Classification

To further resolve the RNA-associated conformational landscape, an additional classification based directly on experimentally derived Rg and COM threshold values separated the ensemble into closed, intermediate, and open states. In this scheme, the closed population corresponded primarily to apo-like conformations, while intermediate states mapped closely to elongation-like structures and the most expanded conformations resembled pre-translocation/open RNA-bound states. These latter states represented the extreme range of thumb–interface opening sampled during the simulations, suggesting a continuum of RNA-accessible conformations rather than a single bound-state geometry.

Together, these classifications establish a structurally interpretable conformational framework linking MD-derived ensembles to experimentally observed functional states. The identification of distinct opening regimes and transitional intermediates further highlights conformational substates that may expose transiently accessible pockets or alter interdomain communication, providing a basis for future exploration of state-selective and allosteric inhibitor design. Furthermore, mapping dominant conformational states may facilitate identification of structurally relevant ensembles associated with ligand accessibility and potential allosteric targeting.

## 3. CONCLUSIONS

How viral RNA-dependent RNA polymerases (RdRp) coordinate local catalysis with global conformational changes remains poorly understood, yet this coordination is essential for replication and represents a potential target for antiviral therapy.^27^ Using microsecond-scale molecular dynamics (MD) simulations of the SARS-CoV-2 nsp12 polymerase, we show that the RdRp-thumb acts as a central dynamic integrator linking active-site readiness, RNA-binding site accessibility, and long-range allosteric communication (**Fig. 9D**).

Our analysis reveals that the thumb subdomain functions as a conserved, mobile gating element whose motions are tightly coupled to global conformational changes, while the coronavirus-specific interface domain acts as a stabilizing scaffold that constrains inter-domain flexibility (**Figs. 4, S5**). This dual architecture suggests the possibility of pursuing two complementary antiviral strategies: targeting the thumb as a dynamic, ligand-accessible regulatory site, and disrupting the interface domain to impair long-range structural coupling required for processive RNA synthesis. *Together, these findings define mechanistically distinct and druggable control points in the coronavirus replication machinery*. Beyond coronaviruses, these findings also support a general model in which thumb-subdomain dynamics govern accessible conformational states of viral RdRps, establishing a mechanistic framework for the development of *broad-spectrum allosteric antivirals* that exploit conserved regulatory features across RNA polymerases.

Notably, our characterization of the apo nsp12 ensemble is dominated by coordinated motions in which changes in global polymerase compaction (radius of gyration, Rg; **Figs. 8-9**) occur together with shifts in thumb–interface separation (center-of-mass, COM; **Figs. 4, S5, 8-9**), indicating that the *thumb subdomain dynamics is structurally coupled to large-scale conformational rearrangements*. Projection of experimentally determined structures onto the Rg–COM free-energy surface enabled assignment of NAC-cycle functional states across the MD-derived conformational landscape, revealing discrete basins corresponding to apo-like (empty) and elongation-like (RNA-bound) configurations (**Fig. 9D**). These states are further organized along a coherent structural axis separating compact and expanded ensembles across experimentally validated conformations (**Fig. 2**, **Fig. 8 legend**). This organization suggests that the apo polymerase samples discrete preorganized conformational substates associated with distinct functional stages of the NAC replication cycle, rather than a continuum of nonspecific structural fluctuations. Integration of RMSD/RMSF (**Figs. 3, S2)**, COM-distance (**Figs. 4, S5**), and DCC analyses (**Fig. 5**) further indicates that these localized thumb subdomain rearrangements propagate throughout the polymerase scaffold, influencing overall structural stability and inter-domain coordination.

Importantly, the distribution of the trajectory ensemble closely recapitulates the Rg–COM features observed for the NAC structures, providing independent support for the conformational assignments (**Fig. 9D)**. The lowest Rg and minimal interface-thumb COM displacement are observed in the apo^2^ and NiRAN-associated Capping^26^ conformations (**Figs. 2, 9D, Table S2**), indicating a globally compact architecture with reduced inter-domain mobility. This ensemble is consistent with a structurally constrained polymerase in which the interface–RdRp organization (**Fig. 4, S5**) stabilizes the core architecture and limits large-scale domain breathing. In contrast, elongation and post-translocation states exhibit intermediate increases in both Rg and COM variability, reflecting a more expanded but still organized structural regime. In these conformations, the RdRp core accommodates RNA duplex propagation while maintaining a relatively stable catalytic geometry.^5,6^ The system thus retains structural integrity while enabling nucleotide addition and template-product translocation. Interestingly, a Remdesivir-stalled polymerase^5^ (**Fig. 9D, Table S2**) exhibits a similar conformational regime, with Rg-COM values closest to those of the active elongating enzyme. The highest values of Rg and COM are observed in the pre-catalytic^19^ and pre-translocation^4^ ensembles (**Figs. 2, 9D, Table S2**). These states correspond to maximal global expansion and increased domain rearrangement, indicating a structurally permissive but dynamically strained regime. This expansion likely reflects the conformational preparation required for nucleotide incorporation and active-site reorganization, consistent with transient opening of the polymerase cleft.

From an allosteric perspective, “druggable” conformations are not necessarily aligned with the most active or most inactive states of the polymerase, but instead reside in intermediate regions of the conformational landscape where structural stability and dynamic flexibility coexist (**Fig. 9D, E-F**). Extreme pre-translocation expansions (**Fig. 9F**, extended ensemble) are too transient for stable pocket formation, while compact NiRAN-associated capping intermediates (**Fig. 9F**, compact ensemble) are limited in allosteric leverage due to restricted inter-domain coupling. In contrast, intermediate states along the transition toward elongation (**Fig. 9F**, intermediate ensemble) provide a structurally stable yet mechanically responsive regime in which perturbation of (sub)domain coupling can efficiently bias the system away from productive catalysis. Within this framework, “conformational trapping” of the thumb subdomain emerges as an ensemble property of the apo polymerase, where subsets of configurations are preorganized into inactive geometries (i.e., stabilization of non-productive states within the free-energy landscape), defining a structurally grounded mechanism that may be exploitable for antiviral intervention.

## 4. MATERIALS and METHODS

### 4.1 Structural model

The structure of the SARS-CoV-2 nsp12 polymerase in complex with cofactors nsp7 and nsp8 was obtained from the RCSB Protein Data Bank (PDB ID: 6M71; 2.90 Å resolution).^2^ The apo complex comprises nsp12 (chain A) with cofactors nsp7 (chain B) and nsp8 (chains C and D), derived from X-ray structures of proteins expressed in *Escherichia coli* BL21 (DE3). The structure was prepared using the Prime package as a prediction tool to fill in missing gaps and residues.^28^ An initial restrained minimization through the Protein Preparation Wizard^29^ was executed to ensure proper structure completion and verify lack of residue clashing. To maintain four separate chains, TER lines were inserted manually in the pdb file as indicated by gaps in tleap.^30^ The system was then solvated in a water-box with a 10.0 Å buffer distance and K^+^ ions were added to neutralize the overall system charge. In addition, 0.15M of KCl was added into the water box to account for the salts present under physiological conditions and to maintain consistency between both systems. The complete solvated apo system has approximately more than 151,000 atoms.

### 4.2 Molecular dynamics (MD) simulations

All-atom molecular dynamics (MD) simulations were performed using the AMBER framework,^31^ using the ff14SB protein forcefield^32^ paired with TIP3P water model^33^. The total simulation time was 1 *μs*. The time step for integration was 2fs for these systems. Langevin dynamics were applied for a consistent temperature of 303 K with a collision frequency of gamma = 1/ps. Periodic boundary conditions were applied under constant pressure with the chosen thermostat. Bond length constraints within the system included in hydrogen bonding were specified via the SHAKE algorithm.^34^ A 10.0 Å cutoff was applied for calculations of Coulombic interactions alongside the Particle Mesh Ewald (PME)^35^ procedure for longer-range electrostatic interactions. Simulations underwent a 10 ns minimization to relax explicit waters, counterions, and apo protein complex after an initial restrained structure minimization was performed in Prime. The Langevin thermostat was maintained for the duration of heating and equilibration with anisotropic Berendsen weak-coupling barostat^36^ for pressure control. Two step heating procedures were carried out keeping the apo protein complex fixed with a force constant of 10.0 *kcal*/*Mol* Å^“^ while heating to 100 K then proceeding to 303 K over 5 ns using constant pressure boundary conditions. The system was held at a constant temperature of 303 K for 6 nanoseconds with fixation constants absent. Simulations including the apo protein complex were performed utilizing the GPU ensemble of AMBER 20.^37,38^

### 4.3 Structural descriptors of conformational dynamics

**Root-mean-square deviation** (*RMSD*(*t*)) of backbone atoms and **root-mean-square fluctuation** ((*RMSF*(*i*))) of Cα atoms were computed using cpptraj^39^ from the AMBER simulation package, with *RMSD*(*t*) evaluated relative to the initial structure.

**Center-of-mass (COM) distances** between selected structural elements were calculated using MDAnalysis.^40,41^ Atom groups corresponding to the regions of interest were defined via MDAnalysis selection syntax, and their center-of-mass (COM) positions were computed for each trajectory frame. The interdomain separation was defined as the Euclidean distance between the COMs of the two groups:

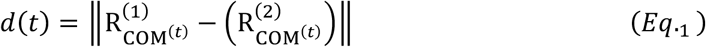

Where 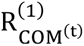 and 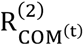 denote the time-dependent COM positions of the selected structural regions. Time series of d(t) were extracted using simulation timestamps and analyzed using NumPy.^42^ Distributions of d(t) were obtained from histograms of sampled values. Raw data were exported for downstream analysis, and all plots were generated using Matplotlib.^43^

**Local structural compactness** was quantified using the radius of gyration, *Rg*(*t*), computed for backbone Cα atoms of selected residues as in Ref. ^5^. *Rg*(*t*) was defined as the mass-weighted root-mean-square distance of atoms from their center of mass and was evaluated for each trajectory frame using MDAnalysis.^40^ All numerical processing was performed using NumPy.^42^

### 4.4. Conformational landscape analysis

#### Dimensionality Reduction by PCA and Clustering

To characterize the conformational landscape sampled during the simulations, dimensionality reduction and clustering analyses were performed on backbone atomic coordinates and *C*_!_ atomic coordinates. Trajectory structures were first aligned to the initial frame to remove global translational and rotational motions using MDTraj.^44^ The resulting Cartesian coordinates were reshaped and projected into a low-dimensional space using principal component analysis (PCA), retaining the dominant collective motions of the system. Conformational states were identified in the reduced principal component space using both density-based (DBSCAN)^45^ and centroid-based (k-means)^24^ clustering approaches. The trajectory was aligned to the initial structure and projected onto the first principal components (PC1–PC3) using atomic coordinates (either backbone or *C*_!_), and time-dependent projections were constructed using simulation timestamps. The fraction of variance captured by each principal component and the cumulative variance were computed to assess convergence of the essential dynamics space. The resulting projections were analyzed as a function of simulation time, and all statistical processing and data handling were performed using NumPy^42^ and Pandas.^46^ Visualizations of variance profiles and conformational sampling in principal component space were generated using Matplotlib.^43^ Density-based clustering was performed using density-based spatial clustering of applications with noise (DBSCAN)^45^, enabling identification of densely populated conformational regions without assuming a predefined number of states. In parallel, centroid-based clustering was performed using the k-means algorithm^24^ to further partition the conformational space and define representative cluster centroids. Both clustering approaches were applied to projections onto the first two principal components using scikit-learn^47^.

#### Collective dynamics analysis

Dominant collective motions of the system were characterized combining PCA and dynamical cross-correlation analysis. Trajectory structures were first aligned to the initial frame to remove global translational and rotational motions using backbone atom selections. Next, PCA eigenvectors and variances were extracted from the trajectory and residue-wise amplitudes along PC1 and PC2 were calculated from the squared Cartesian displacements of each Cα atom. Normalized per-residue amplitudes were then used to identify structural regions contributing most strongly to the collective modes and were mapped onto the protein structure (e.g., through B-factor encoding for visualization). In parallel, dynamical cross-correlation maps (DCCM) were computed to quantify correlated atomic fluctuations over the simulation trajectory. Positional fluctuations were extracted from the aligned trajectory and used to construct a residue-wise cross-correlation matrix describing pairwise dynamic relationships between atomic displacements. The resulting matrix was analyzed to identify regions of correlated and anticorrelated motion. All numerical processing and matrix operations were performed using NumPy^42^, while visualization of principal component projections, variance profiles, and correlation matrices was carried out using Matplotlib^43^. [PCA was used both to define conformational states (landscape analysis) and to characterize dominant collective motions (dynamical analysis), serving distinct but complementary purposes.]

### 4.5. Free energy landscape reconstruction

#### Kernel density estimate (KDE) & free energy surface (FES)

The conformational free energy landscape was reconstructed from the joint distribution of radius of gyration, *Rg*(*t*), and center-of-mass (COM) displacement, *d*(*t*), obtained from MD trajectories. A two-dimensional kernel density estimate (KDE) was computed over the (*Rg*(*t*), *d*(*t*)) space using a Gaussian kernel to approximate the configurational probability density *P*(*Rg*, *d*). The free energy surface (FES) was obtained via Boltzmann inversion:^48–50^

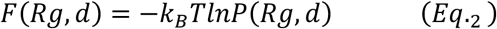

where *k*_)_is the Boltzmann constant and *T* is the simulation temperature (303 K). The resulting free energy landscape was shifted such that the global minimum was set to zero for visualization. The resulting landscape was used to identify metastable basins and transition regions sampled during the simulations. One-dimensional free energy profiles were additionally computed by marginalizing the KDE along individual collective variables, *Rg*(*t*) and *d*(*t*), providing reduced representations of conformational stability and state populations.

To assess the effective dimensionality of the sampled conformational space, PCA was performed on the joint (*Rg*(*t*), *d*(*t*)) dataset. Each trajectory frame was represented as a two-dimensional vector (*Rg*(*t*), *d*(*t*)), which was mean-centered prior to construction of the covariance matrix:

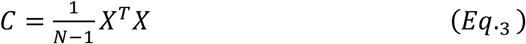

where *X* is the centered data matrix and *N* is the number of frames. Eigenvectors of *C* define the principal components, and eigenvalues quantify the variance captured along each mode. The relative variance explained by the principal components (94.94% and 5.06% for PC1 and PC2, respectively) was used to evaluate whether the conformational landscape is dominated by a single collective coordinate or distributed across multiple independent degrees of freedom.

#### Conformational state decomposition & experimental NAC mapping

Conformational space was analyzed by projecting MD trajectories onto the radius of gyration (Rg) of RNA-binding residues and the thumb–interface center-of-mass (COM) distance.

#### Hexbin density plot

Conformational sampling in Rg–COM space was visualized using hexagonal binning implemented in Matplotlib ^43^(plt.hexbin). Rg and thumb–interface COM distances were projected onto a two-dimensional state space, discretized into hexagonal bins (gridsize = 35). Bin color intensity reflects the number of trajectory frames within each bin, enabling visualization of densely populated conformational regions while reducing overplotting associated with conventional scatter plots. NAC and other representative experimental structures were mapped onto the hexbin density plot and are shown as dots (for RNA bound) or stars (empty structures), colored according to the legend in Fig. 2.

#### Global state decomposition

Unsupervised clustering was performed using k-means^24^ (k = 2), yielding two dominant conformational populations. Cluster centroids were mapped onto experimentally derived nucleic acid cycle (NAC) representative structures in the same Rg–COM space to assign functional labels. This mapping identified correspondence with apo and elongation states, which were used to designate “empty-like” and “RNA-bound-like” ensembles, respectively.

#### Finer state resolution

Structures were additionally classified into Compact, Intermediate, and Extended populations based on threshold-based segmentation along the COM coordinate (57.0 and 58.0 Å; Table S2). Mean Rg and COM values were computed for each state to define representative configurations. Agreement with experimental NAC structures was assessed by comparing state-averaged coordinates with reference apo and elongation conformations.

All images were rendered in Maestro (Schrödinger, LLC, New York, NY, 2021.), ChimeraX^51^ or VMD.^52^

## Author Contributions

HGG performed the molecular simulations, analyzed the data, and contributed to manuscript writing. AB analyzed the data. PBM contributed to supervision of the computational studies, data analysis, and manuscript writing. EG supervised the computational studies, conceived the research, and wrote the manuscript. All authors critically reviewed and edited the manuscript.

## Funding Sources

This research was supported by The RF Research Foundation and the City University of New York (CUNY) Queens College startup funds (EG), the PSC-CUNY Award # 67118-00 55 (EG) and the CUNY Faculty Fellowship Publication Program (FFPP) Office of Faculty Affairs, The City University of New York (EG), Saint Joseph’s University (SJU), and the West Center for Computational Chemistry and Drug Design at SJU.

## Competing Interests

The authors declare no competing interests.

## Data Availability Statement

Atomic coordinates of representative computational models for the empty and RNA-bound states, together with compact, intermediate, and extended conformational states sampled across the ensemble, are available from the authors upon reasonable request.

## ACKNOWLEDGMENT

Artificial intelligence tools (e.g., ChatGPT OpenAI) were used to assist with literature searches and to improve grammar and manuscript readability. All scientific content, interpretations, and conclusions were reviewed and validated by the authors.

## ABBREVIATIONS

SARS-CoV-2: severe acute respiratory syndrome coronavirus 2
CoV: coronavirus
FES: Free energy surface
KDE: kernel density estimate
NAC: nucleic acid cycle
PCA: principal component analysis
RTC: replication and transcription complex.

## Notes

### Competing Interest Statement

The authors have declared no competing interest.

