## Supporting Information for "Preorganized RdRp-Thumb Dynamics Drives SARS-CoV-2 Polymerase Function"

### Table of content

**Supplementary Figures** 3

**Supplementary Tables** 11

### Supporting Information

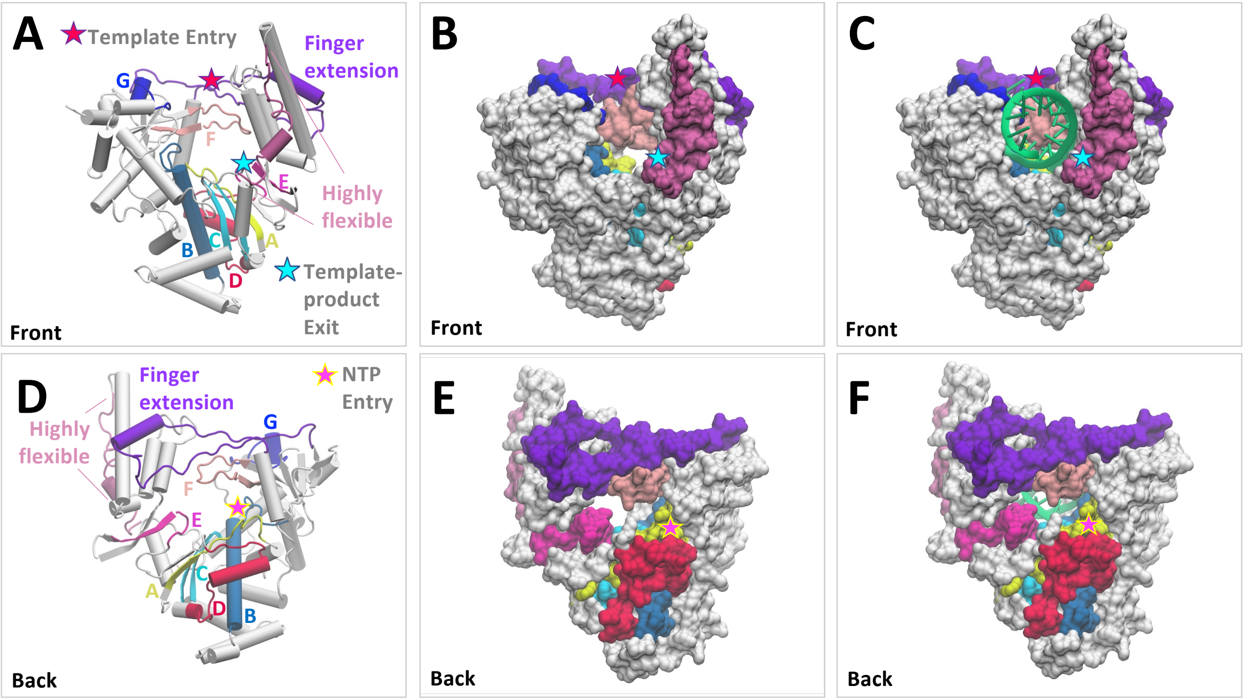


**Figure S1. Structure and RNA channel topology of the SARS-CoV-2 nsp12 RdRp domain.** 3D structure representation of nsp12 RdRp, front (A)-(C) and back (D)-(F) views. Structural motifs, RNA template entry and template-product exit channel highlighted. The polymerase motifs are shown as follows: motif A in yellow; motif B in pale blue; motif C in cyan; motif D in red; motif E in mauve; motif F in pink; motif G in blue. Residue numbers are attributed as indicated in **Figs. S1** and **S2**. The finger extension region (amino acids 401-447) is shown in purple. The highly flexible region (amino acids 907-929) in the thumb subdomain is highlighted. Protein atoms are shown in tube and surface representation in (A)-(D), and in (B)-(C) and (E)-(F), respectively. In (C) and (F), RNA duplex is shown, colored in green (both primer and template strands).

**Supporting Information**

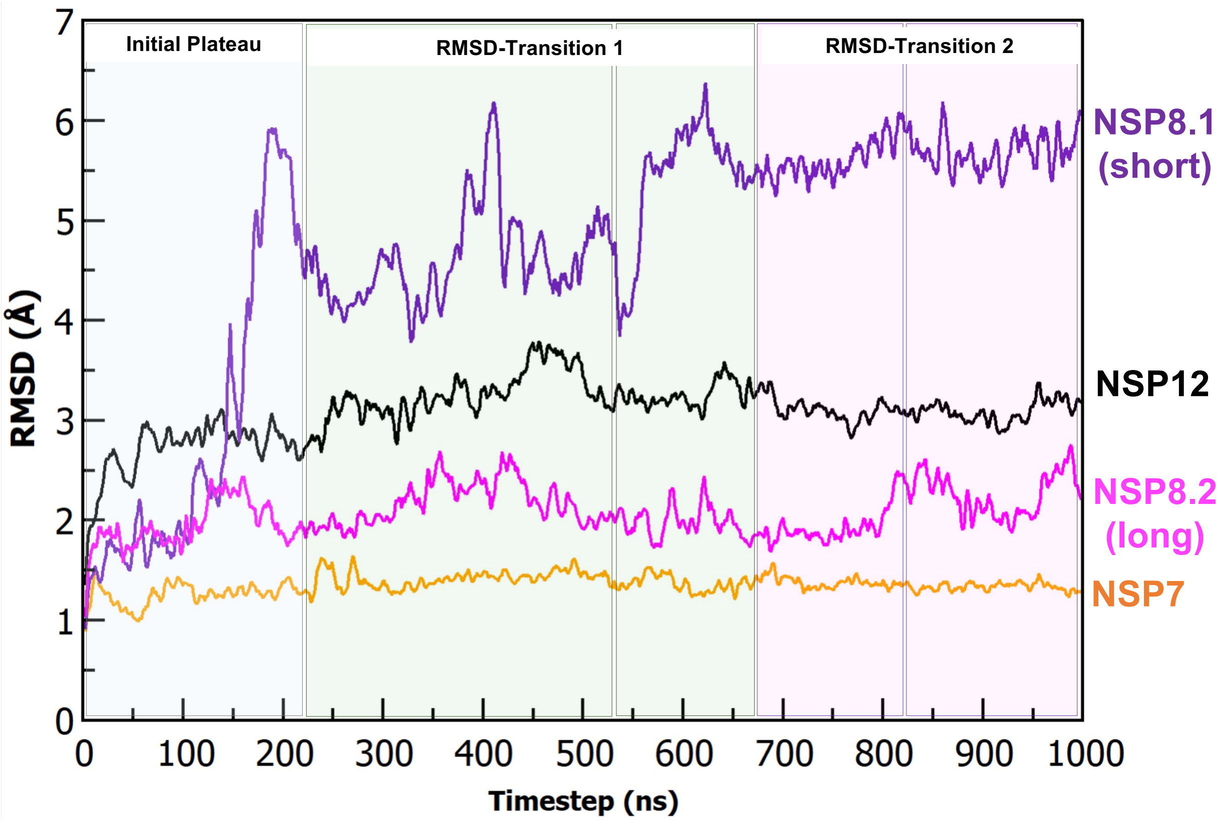


**Figure S2. RMSD of apo SARS-CoV-2 NSP12 polymerase and cofactors.** Root-Mean-Squared-Deviation (RMSD) of the full length nsp12 polymerase with relative cofactors nsp8 (two protomers) and nsp7 depicts overall conformational deviation of each nsp from their initial starting conformation over time. One nanosecond is equivalent to one conformation. Running average of 5 was used for all graphs for visualizing smoothed trends of the data. Comparison with domain RMSD (Fig. 3B) is highlighted in color.

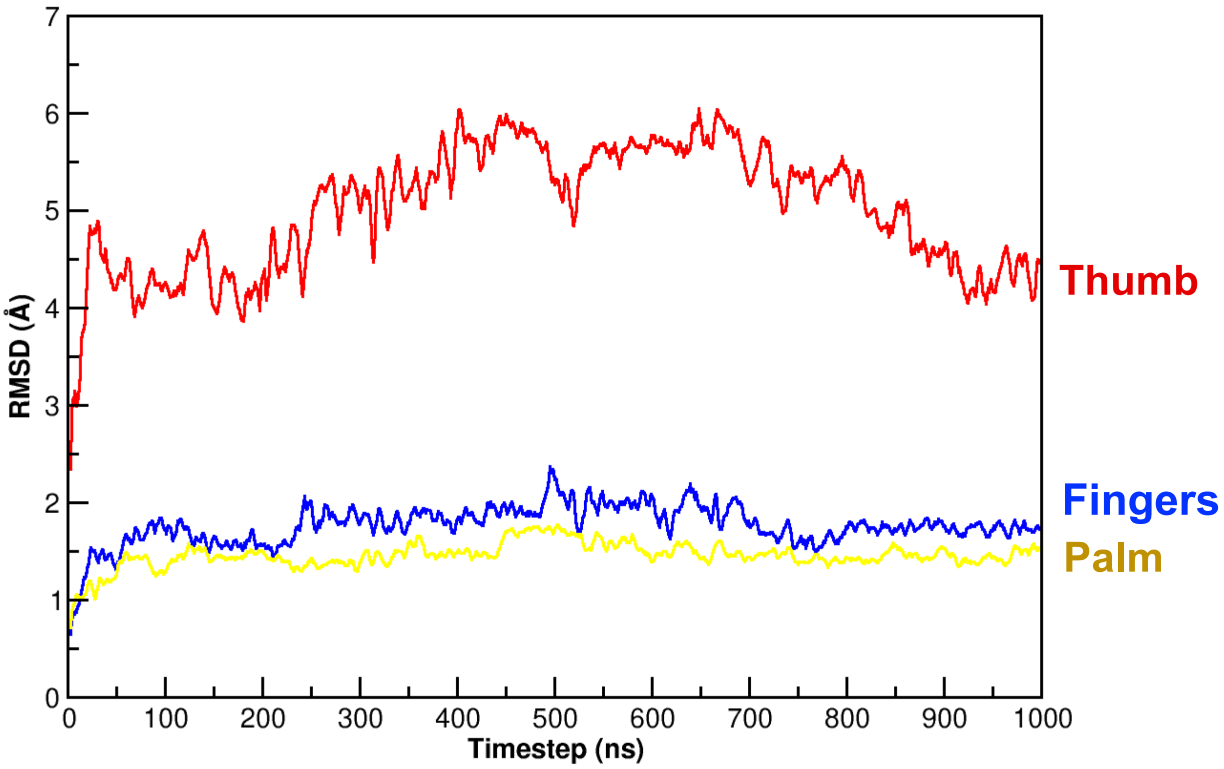


**Figure S3. RMSD of apo SARS-CoV-2 RdRp subdomains.** Root-Mean-Squared-Deviation (RMSD) of the fingers, palm and thumb subdomains of the nsp12 RdRp polymerase. The plot depicts overall conformational deviation of each RdRp domain from their initial starting conformation over time. One nanosecond is equivalent to one conformation. Running average of 5 was used for all graphs for visualizing smoothed trends of the data.

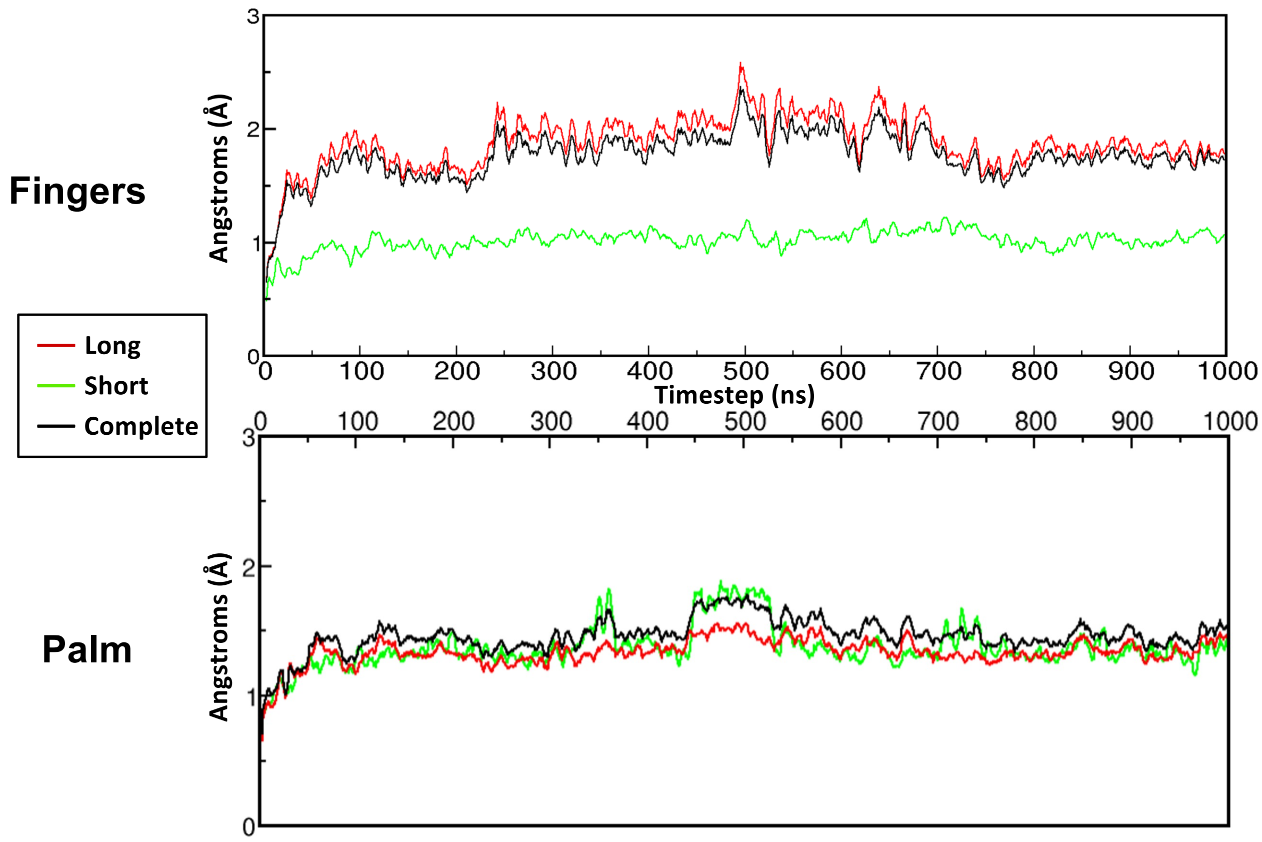


**Figure S4. Split RMSD of SARS-CoV-2 fingers and palm.** Split Root-Mean-Squared-Deviation (RMSD) of the fingers and palm subdomains of the nsp12 RdRp polymerase into their sub-components (long and short), as shown in Fig. 1, compared to the full domain (complete). The plots depict overall conformational deviation of each region from their initial starting conformation over time. One nanosecond is equivalent to one conformation. Running average of 5 was used for all graphs for visualizing smoothed trends of the data.

### Supporting Information

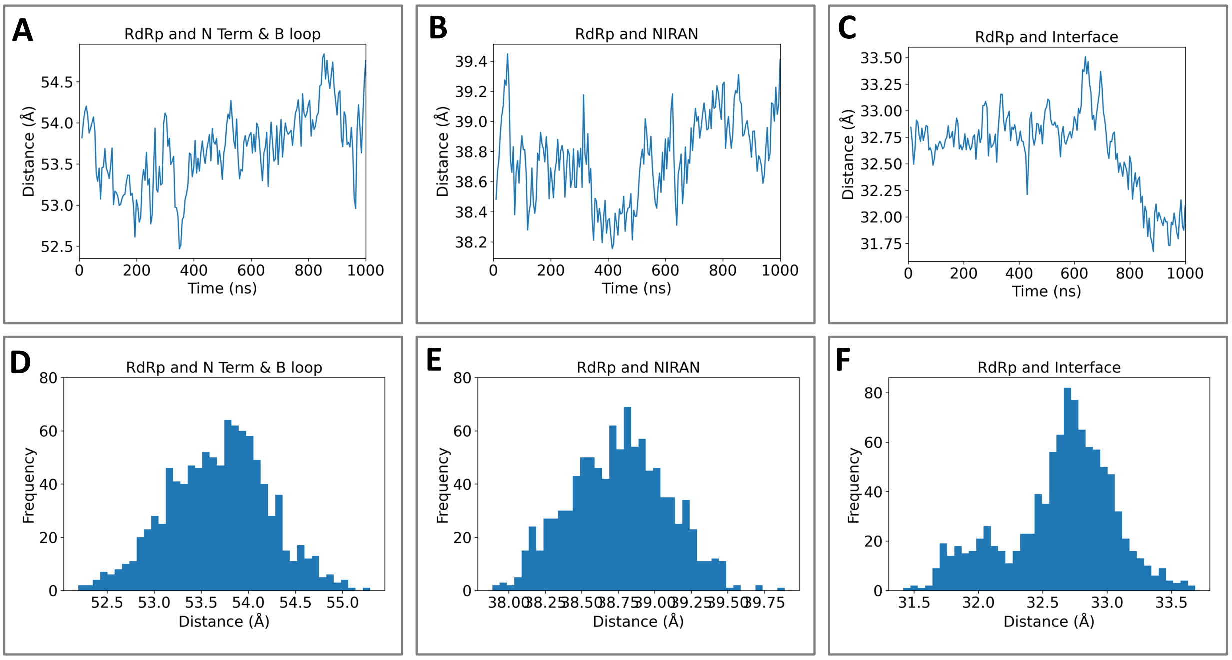


**Figure S5. Center-of-mass (COM) distance analysis of inter-domain organization relative to the RdRp core in nsp12.** (A)-(C) Time series of COM distances between the RdRp core and individual domains (interface, NiRAN and N-terminal with $\beta$-hairpin) over the MD trajectory (1 $\mu$s), reporting large-scale rearrangements of peripheral domains with respect to the catalytic core. Distances exhibit time-dependent fluctuations, with the largest variability observed for RdRp and the N-terminal domain with $\beta$-hairpin, and pronounced variability observed for thumb–interface separations, while RdRp–NiRAN distances remain comparatively constrained. (D)-(E) Probability distributions (histograms) of the corresponding COM distances, quantifying the extent and heterogeneity of conformational sampling. Broad and, in some cases, multimodal distributions for RdRp–interface distances indicate the presence of multiple configurational states, whereas narrower distributions (for RdRp–NiRAN and N-terminal region with hairpin) distances are consistent with a more stable spatial arrangement. Overall, comparative representation of COM distance populations highlighting the relative positioning of domains with respect to the RdRp core. Differences between distributions reveal heterogeneous sampling of inter-domain configurations and define preferred spatial arrangements associated with distinct conformational states of the polymerase.

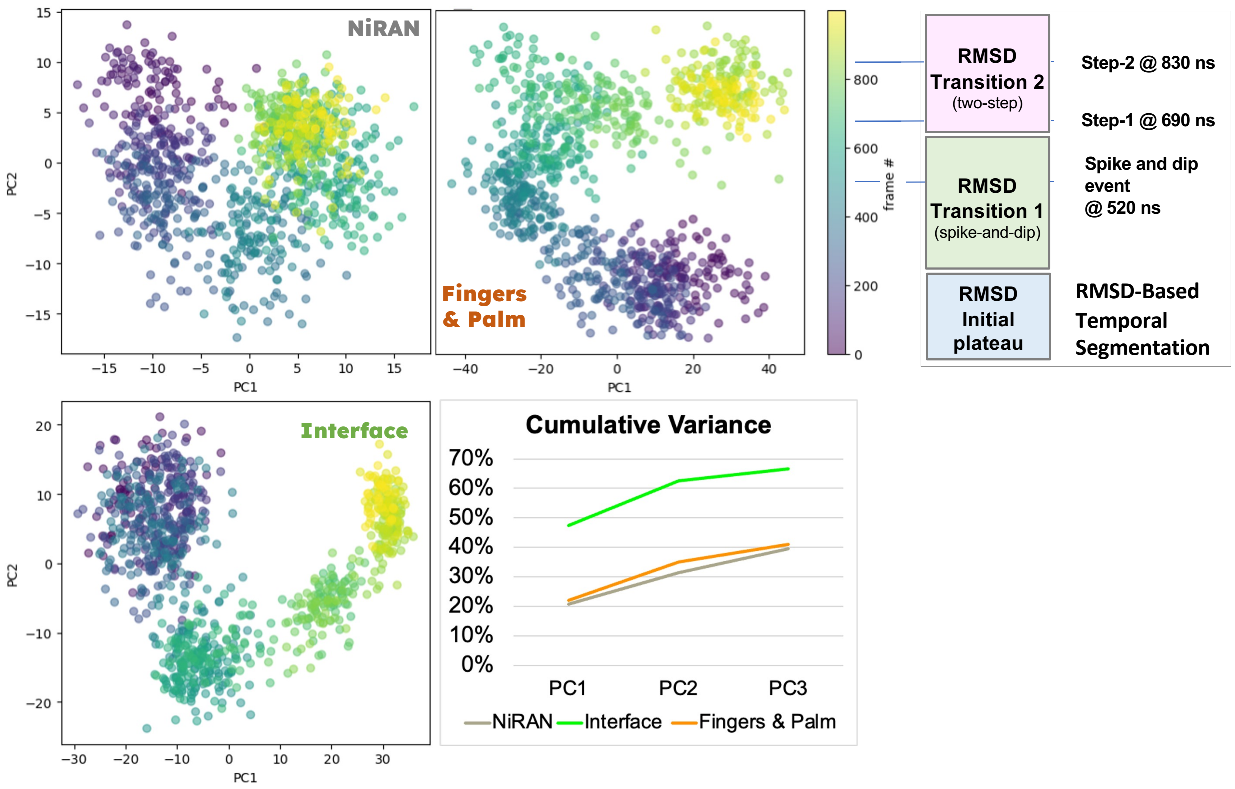


**Figure S6. Principle Components Analysis (PCA) and Cumulative Variance of apo nsp12 NiRAN, Fingers and Palm, and Interface (Sub)Domains.** The PCA graphs, labeled for each structure analyzed with color scheme from **Figure 1**, shows the transformed data using two principal components for each. These uncorrelated points are colored by frame number using the gradient at the right of the graphs (each frame corresponds to 1 ns, for a total of 1000 frames or 1 $\mu$s). The fourth plot at the bottom right is the cumulative variance for each analyzed structure overlaid to demonstrate the enrichment as the structure analyzed becomes specific. RMSD-based temporal segmentation is also shown for comparison with PCA projections (top right). Schematic view of major conformational regimes identified from the RMSD trajectory, included to provide temporal context for the PCA analysis in which frames are colored by simulation time. The system initially resides in a stable initial plateau, followed by Transition 1, characterized by a spike-and-dip event at ~520 ns. A subsequent two-step transition (Transition 2) occurs, with changes at ~690 ns (Step 1) and ~830 ns (Step 2). The vertical axis indicates simulation time (frame number). This scheme enables direct mapping of RMSD-defined transitions onto the time-colored PCA projections, facilitating interpretation of how large-scale structural rearrangements are distributed in principal component space.

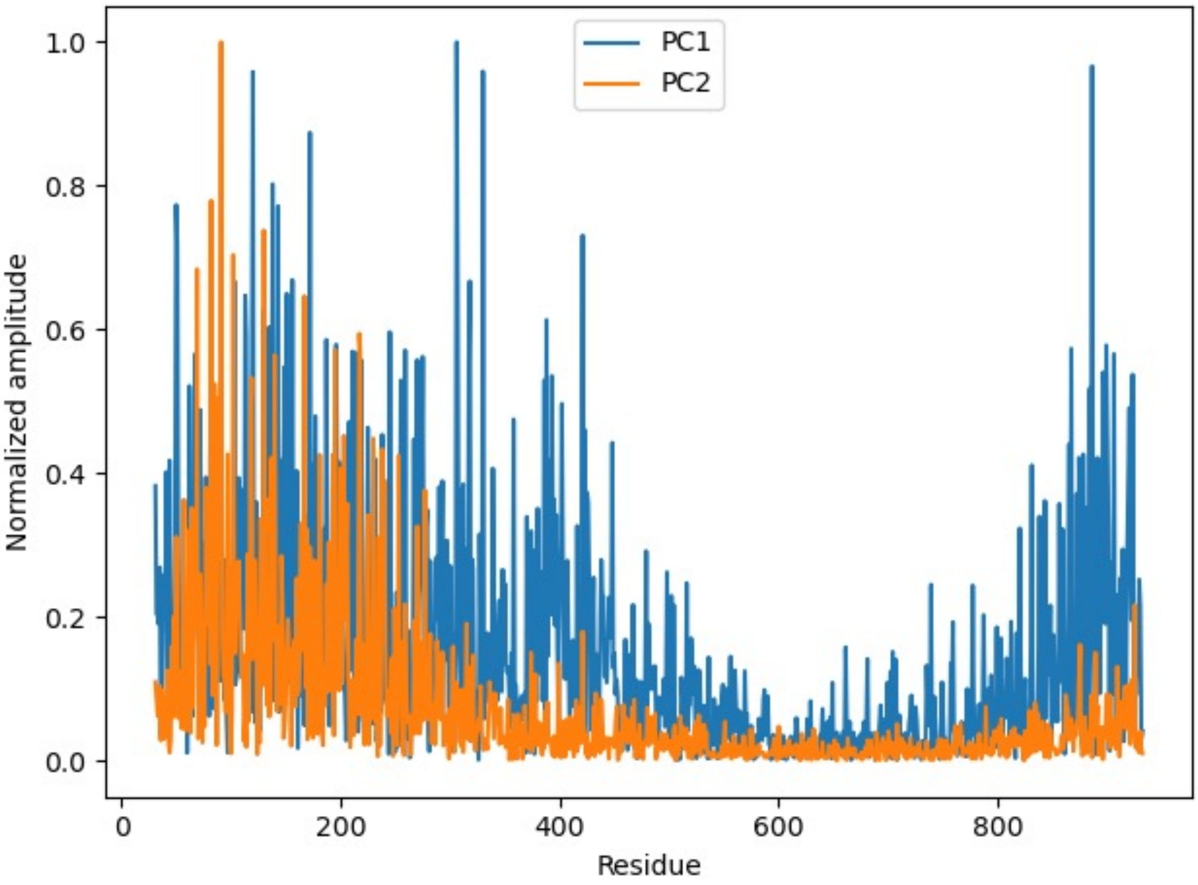


**Figure S7. Per-residue contributions to principal components.** Normalized amplitudes of residue-wise (nsp12 Cα-atoms) contributions to the first (PC1, blue) and second (PC2, orange) principal components derived from PCA of the trajectory. PC1 exhibits distributed contributions across the sequence, with prominent peaks in the N-terminal/NiRAN/interface regions and a secondary increase toward the thumb subdomain, consistent with large-scale coordinated motions. In contrast, PC2 shows more localized contributions, primarily confined to the N-terminal region, with minimal involvement of the polymerase core. Residues within the central RdRp domain display comparatively low amplitudes in both components, indicating limited participation in the dominant collective motions. These profiles identify the regions that drive the principal dynamical modes of the system.

### Supporting Information

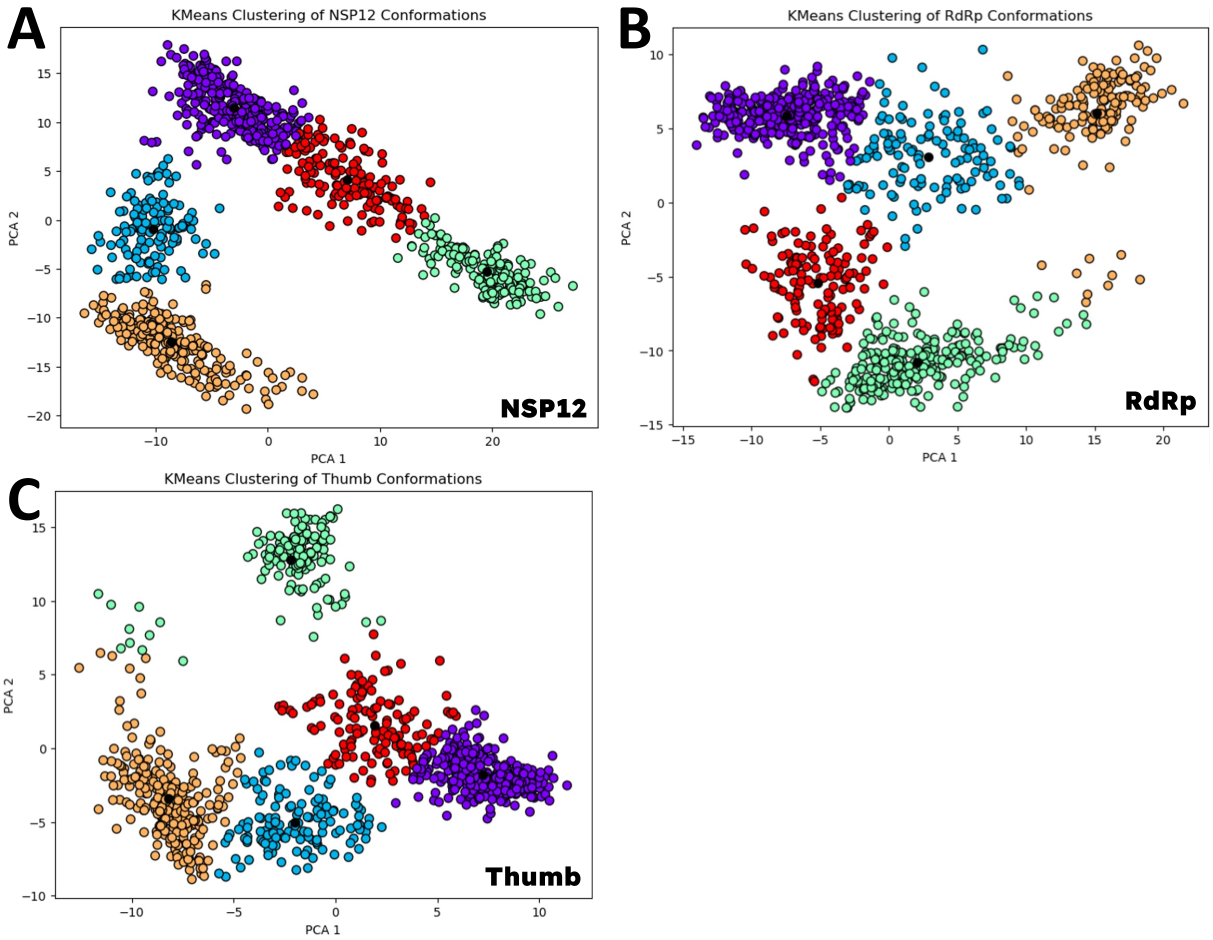


**Figure S8. K-means clustering of Apo NSP12.** Apo nsp12, RNA-dependent RNA polymerase (RdRp) and the isolated thumb subdomain are shown (A to C, respectively). K-means clustering applied to each system partitions the conformational ensemble into *k* groups of structurally similar states. Points are colored by cluster membership, and black markers denote cluster centroids (representative structures).

### Supporting Information

#### Supplementary Tables

**Table S1. Principal component analysis (PCA) of full-length nsp12 and domain-resolved conformational dynamics.**

| Protein, Domain or Subdomain | PC1 | PC2 | PC3 |
|---|---|---|---|
| NSP12 | 24% | 42% | 49% |
| RdRp | 26% | 56% | 52% |
| Thumb | 31% | 58% | 66% |
| NiRAN | 21% | 32% | 40% |
| Interface | 48% | 63% | 67% |
| Fingers & Palm | 22% | 35% | 41% |

*Variance explained by the leading principal components (PCs) obtained from PCA of the MD trajectories for full-length nsp12 and selected structural (sub)domains, including the RdRp core domain, the thumb subdomain, the interface and NiRAN domains. Reported values correspond to the percentage of total conformational variance captured by each principal component.*

**Table S2. Structural descriptors of NAC representative conformations and experimentally resolved reference states of the SARS-CoV-2 nsp12-RdRp complex.**

| State | PDB ID(s) | COM (Å) | Rg (Å) |
|---|---|---|---|
| Apo RdRp* | 6M71 | 56.95 | 15.5 |
| PreTranslocation | 7C2K | 58.86 | 16.04 |
| PostTranslocation | 7BZF | 57.72 | 15.94 |
| RDV-Stalled | 7BV2 | 57.44 | 15.84 |
| Backtracked | 7KRN | 58.29 | 15.78 |
| NiRAN Capping* | 7THM | 56.79 | 15.4 |
| Elongation | 7B3D | 57.49 | 15.76 |
| PreCatalytic | 6YYT | 58.80 | 15.82 |

*NAC representative states and experimentally resolved conformations used for comparison in the conformational landscape analysis. Values include the corresponding Protein Data Bank (PDB) identifier, thumb-interface center-of-mass (COM) distance, and radius of gyration (Rg) of RNA binding residues, both reported in Å. NAC representatives correspond to conformations identified from the experimental ensemble and mapped onto the MD free energy landscape.*
*Entries marked with an asterisk (*) denote apo/empty conformations lacking bound RNA, whereas all remaining structures correspond to RNA-bound replication states.*
